# Ecological drift in the flour beetle microbiome and associated instability in host benefits

**DOI:** 10.1101/2025.09.16.676705

**Authors:** Pratibha Sanjenbam, Harshith Koppa Guruswamy, Rittik Deb, Shivansh Singhal, Sneha Garge, Shyamsunder Buddh, A Dhanush, Al Maman Haque, Shubha Govindarajan, Deepa Agashe

## Abstract

Many eukaryotes harbor complex microbial communities that provide significant benefits to the host. However, recent work demonstrates substantial variation in microbiota across hosts and over time, though the causes and consequences of this instability are not always clear. We describe large temporal, among-individual and across-population variation in the microbiome of the red flour beetle *Tribolium castaneum*, a widespread pest of stored grain flour and a laboratory model organism. Across 5 years, the total load and composition of the bacterial community of laboratory beetle populations fluctuated dramatically, cycling between a few dominant taxa and hundreds of rare taxa. We show that this ecological drift is caused by a lack of maternal microbial transmission and large microbial load fluctuations in flour, which is the primary reservoir of microbial inocula. Newly hatched larvae (from eggs) and newly eclosed adults (from pupae) acquire microbes from ingested flour containing conspecific feces. The flour microbiome fluctuates with host population density and stage structure, with larval feces increasing microbial load and antimicrobial secretions from adults decreasing it. Periodic flour replacement during population maintenance also dramatically decreases flour bacterial load, contributing to stochasticity in microbial colonization. The resulting drift was associated with unstable fitness benefits for beetles, though we cannot disentangle cause vs. effect. Our work thus represents a case study of significant ecological drift even in a highly controlled laboratory environment. We suggest that such unstable microbial communities and variable host benefits may not be uncommon, and offer an opportunity to analyze early stages of the evolution and ecology of host-microbial interactions.

## INTRODUCTION

Animal-microbe relationships are seemingly ubiquitous, and thus the focus of a large body of biological research, especially on obligately host-associated microbes e.g., see (1,2). However, several mammals, insects, and birds host transient or unstable microbiomes, and/or do not appear to acquire any specific benefits from the microbes that they harbour e.g., (3–7). Many host species also show high individual variation in their microbiomes (reviewed in, e.g., (8), even across individuals reared in identical conditions in the laboratory (9–11). What mechanisms and processes drive this variation?

Variability in the microbiota of a host species suggests a strong role for stochastic processes in driving microbial community assembly and maintenance. However, these processes may operate across different scales of organization, including at the level of the meta-community (8). Broadly, variation may arise from historical contingency or limited dispersal, or it may reflect ecological drift, whereby stochastic processes (birth, death, and dispersal) rather than deterministic processes govern the abundance of specific microbes (12). For instance, Vega and colleagues inoculated *Cenorhabditis* worms with similar numbers of two competitively equivalent *Escherichia coli* strains (10). At high inoculum densities, most worms were reliably colonized by a nearly equal mixture of both strains. But at low inoculum density, demographic stochasticity determined the outcome, with most worms colonized by only one of the two strains. Such ecological drift should be stronger when unopposed (or weakly opposed) by selection favouring obligate, consistent mutualism; for example, when hosts do not gain significant fitness advantages by harbouring specific microbiota. On the other hand, determinism in community assembly does not necessarily reflect strong selection, because other processes — such as a strong environmental influence coupled with stable environments, or dietary specialization — may also drive deterministic colonization and assembly dynamics. An interaction between host- and microbe-specific factors may also lead to between-host microbiota variation; e.g., a combination of host control and microbial colonization may drive distinct routes of microbiome assembly in different individuals. Thus, several factors and processes may combine to drive microbiome variation across hosts. For example, strains of the symbiont *Lactobacillus plantarum* isolated from wild *Drosophila melanogaster* showed positive density-dependent colonization (i.e., were strong colonizers) — as observed in worms, discussed above — but strains isolated from laboratory flies or humans remained consistently poor colonizers regardless of density (9). This study also uncovered additional effects of stochastic population bottlenecks, presence of competitor strains, and host genetics, highlighting the complexity of mechanisms underlying host-to-host microbiome variation. Even in a neutral model of community assembly (i.e., under no selection), weak dispersal from the environment to the host, coupled with a short host lifespan, led to different microbiome compositions in different hosts (13). Thus, ecological drift may arise from several sources, driving large variation in host microbiomes. However, the relative contribution and consequences of these factors for ecological drift, especially for complex microbial communities, remains unclear.

Here, we focus on host-bacterial interactions in the red flour beetle *Tribolium castaneum*. These beetles are generalist, widespread pests of cereal grain flours, and complete their entire life cycle (egg to larva to pupa to adult) in flour. Hence, the beetles effectively live in a largely closed system, which can be efficiently reproduced in the laboratory under controlled environmental conditions such as temperature and humidity. Prior studies report somewhat diverse sets of flour beetle microbiomes, though generally dominated by bacteria from the family *Enterobacteriaceae*, with some variation across life stages (14–17). Importantly, our previous work indicated a significant (but non-essential) beneficial effect of flour-associated microbes on beetle fitness parameters such as reproduction, survival, and immune function (16,17). However, across these two studies, we observed substantial variation in beetle microbiomes, which was also reflected in results from a different group finding variation in beetle microbiomes across sampling locations (15). Given the observed fitness benefits of the microbiome for beetles, this variation is surprising, and indicated that the non-obligate host-bacterial interactions may be associated with significant ecological drift. Hence, this system presented a unique opportunity to measure and understand the factors responsible for ecological drift.

We first systematically quantified individual and population level as well as temporal variation in the flour beetle microbiome across several years, using previously stored beetle samples from populations that had been maintained in near-identical laboratory conditions. Finding large variation in all these cases, we conducted a series of experiments to determine the causes of the apparent ecological drift. Specifically, we tested how the nature and magnitude of variation in the beetle microbiome was affected by sources of bacterial inocula entering the flour habitat (fresh flour, experimenter identity, and beetle feces) and factors influencing bacterial survival and load in the habitat (beetle-secreted antimicrobials and flour replacement during population maintenance). Finally, we found that beetles do not consistently derive large benefits from their microbiota, potentially because the microbiome is unstable. Thus, we conclude that ecological drift in the flour beetle microbiome is driven by a combination of factors that alter the fecal bacterial inoculum in the flour, as well as the timing of flour change implemented during population maintenance (whereby used flour is replaced with fresh flour). We suggest that similar factors at play in the beetles’ natural habitat (flour and grain warehouses) may help explain why a strong, obligate host-bacterial mutualism has not evolved in this system, despite potential benefits for the host.

## METHODS

### Population maintenance and sample storage

For all our experiments, we used a large outbred stock population, described earlier (17). In brief, we allow 2000-3000 adults (∼2-4 weeks old) to oviposit on 750 g fresh wheat flour for 7 days in a large plastic box, and then remove the adults. After allowing 2-3 weeks for offspring development, we conduct maintenance, replacing the used flour with fresh wheat flour and removing dead individuals and larval molts. At this stage, most individuals are either large larvae or pupae, and these are allowed to develop further for another 2-3 weeks until we see a large number of adults. We then start a new stock generation with a subset of these adults. Three independent replicate boxes of the outbred line are maintained in this fashion, but we used one of them as our focal stock population, and all experiments were done using individuals derived from this population, except where noted otherwise.

Every packet of wheat flour used for our experiments is first frozen at −80°C for 4 hrs to remove any prior pest infestation (17) and is allowed to return to room temperature before use. All beetle populations are maintained in plastic boxes with holes in the lid for ventilation (except when noted otherwise), and kept in dark incubators at 33±1 °C. Beetles or flour samples used for microbiome analysis were stored at −80°C.

### Collecting samples to estimate microbiome variation across time, replicate populations, and generations

#### Experiment 1a

To determine change in beetle microbiome composition in the same lineage over time, in the year 2022, we created a new, parallel stock population. To obtain adults that were exactly 2 weeks old, at each generation we allowed adults to lay eggs only for 48 hours (instead of 7 days in the usual protocol described above) (Fig S1A). Every 3 months, we stored 2 week-old female beetles (n = 10) from the population.

#### Experiment 1b

To expand the above time series, we added samples from the same outbred stock population, which were stored for other projects from 2015–2021. These represent opportunistic samples and are therefore not spaced out systematically across the years. Since these were sampled directly from the stock boxes, precise adult age was not known; however, given the beetle life cycle, we estimated that the females were between 2-4 weeks old, comparable to our 2022 samples. When possible (July 2020, February 2021, and March 2022), we also analysed males from the same population (n=10) to assess sex-specific variation.

#### Experiment 1c

To determine microbiome variation across replicate populations with similar original ancestry, we sampled and stored individuals from the three independent stock boxes described above, in February 2016 and again in March 2022. As described for Experiment 1a, samples for March 2022 were collected following 48 hours of oviposition, and therefore, females were 2 weeks old. In contrast, samples stored in February 2016 were 2-4 week old females (Fig S1B). When available, we also analysed males from the same boxes.

#### Experiment 2

To test for differences between parent and offspring microbiomes, we divided ∼600 2-week-old adults from the stock population into groups of 200, and allowed each group to oviposit in 200 g fresh wheat flour for 7 days, creating three replicate boxes (Fig S1C). This protocol mimics our stock population maintenance. From each replicate box, we sampled 1-week old offspring (n = 10 females and 10 males) for microbiome analysis. These data allowed us to test for (a) variation in parent vs. offspring microbiomes and (b) variation across replicate boxes of offspring of parents from the same stock box.

### Testing the effect of external factors that may influence the beetle microbiome

#### Experiment 3

To test the effect of factors such as experimenter identity, exposure to sterile vs. normal air, and flour sterilization, we allowed 1500 ∼4 week-old adults from the stock population to oviposit for 48 hrs in 600 g double sifted fresh wheat flour (Fig S2A). To measure the effects of air exposure, we sifted out 100 eggs per treatment and placed each set of eggs in 100 g fresh wheat flour, with normal air supply (box lid had holes) vs. sterile air (box without holes, but opened under sterile air supply in a laminar airflow twice a week). Similarly, we generated two more batches of eggs and placed them in 100 g of either normal or dry-heated wheat flour (90°C for 45 mins). In all cases, we allowed eggs to develop for 5 weeks, i.e., until they were 2-week-old adults, and stored for microbiome analysis (n = 5 males and 5 females/treatment). The sterilized wheat flour served as our positive control, wherein we expected a distinct microbiome and lower total bacterial load due to the heat treatment. To test the effect of experimenter identity, we sampled 1500 adults from the stock population and split them equally between two experimenters. Each experimenter allowed adults to oviposit for 7 days in independent boxes containing 600 g double sifted fresh wheat flour. After 5 weeks, the resulting adult offspring (1000 each) served as each experimenter’s independent set of parents. These were allowed to oviposit for 48 hrs in 300 g of double sifted fresh wheat flour, and offspring were sampled when they turned 2-week-old adults.

### Testing for maternal transmission of the beetle microbiome

#### Experiment 4a

To test for maternal microbiome transmission, we allowed ∼50 2-week-old adult beetles to oviposit for 48 hrs on 5 g fresh wheat flour in each of three replicate boxes. We sampled the parents (n = 5 females/replicate) and eggs (n = 3 replicates, with 50 pooled eggs/replicate), and stored them for microbiome analysis (Fig S3A).

#### Experiment 4b

In experiment 4a, contamination of the egg surface from flour or adult feces was a concern. To address this, we directly dissected ovaries and eggs from ∼2 week old stock females (n=3) that were previously stored at −80°C. We rinsed each female in 70% ethanol and washed with sterile saline (0.9% sodium chloride). We placed the beetle on a 5% agar plate and cut the posterior tip of the abdomen using fine scissors (Fine Science Tools) under a stereomicroscope (Leica V4.12). We applied pressure to the sternum to force the abdominal contents outside, then teased apart the ovary with needles and forceps and collected eggs and ovaries from each female for DNA extraction (Fig S3B). We ensured that we did not disrupt the gut during dissection.

#### Experiment 4c

In the above experiment, we used ∼2 week old females. In subsequent experiments (see below), we observed that the adult microbiome took ∼4 weeks to stabilize, and it was possible that vertical transmission could occur at that stage. Hence, we analysed the microbiomes of dissected eggs from females collected from stock populations at 2, 4, and 6 weeks post eclosion (Fig S3C). We dissected them as described above, and collected 3 groups of 10 eggs per age category, with each group containing eggs from 2-5 females.

### Testing the effect of fecal inoculum on the beetle microbiome

#### Experiment 5

To test whether beetle feces serve as a source of bacterial inoculum, we conducted a series of experiments (Experiments 5-8). First, if adult feces are a key source of inoculum, we expected that bacterial load would be positively correlated with adult density. To test this, we took 2-4 week old adults, and created two densities by placing 200 (low) or 2000 adults (high) in 300 g wheat flour. We allowed adults to oviposit for 7 days and then removed them. We split the flour (including eggs) into three replicate lines (100 g each), and allowed offspring to develop for 5 weeks, i.e., until they were 2-week-old adults. We stored offspring at −80°C for microbiome analysis (n = 5 females/replicate/treatment). For one replicate per treatment, we also stored 5 males (Fig S2B).

#### Experiment 6

Next, determined whether the larval and adult fecal microbiomes were different, and changed with age. To obtain individuals of known age, we allowed ∼200 2-4-week-old adults from the stock population to oviposit in 200 g of fresh wheat flour, and removed the adults after 48 hrs. Once we started observing adult offspring (after 4 weeks), we sampled adults every week for 6 weeks, placing them in empty 1.5 ml tubes (Eppendorf) for 24 hrs to collect their feces (n = 30 adults/tube, with 3 replicate tubes per time point). After this, we placed the adults back in their box to maintain the same population density until the next sampling point. Thus, we obtained adult fecal microbiomes from 1-6 weeks of age. Similarly, we obtained larval fecal microbiomes, sampling every 3 days for 21 days, until the larvae began pupating (n = 30 larvae/tube, with 3 replicate tubes per time point). We immediately processed all fecal samples for microbiome analysis, as described in later sections (Fig S4A).

#### Experiment 7

To determine the effect of larval vs. adult fecal inoculum on recipient microbiomes, we added feces to fresh wheat flour and monitored the microbiome of recipient pupae until they were 6-week-old adults (i.e., for 7 weeks total). We first prepared ‘enriched flour’ containing either larval or adult feces, by adding either 1000 larvae (10-14 days old) or 200 adults (2-4 weeks old) to 50 g of wheat flour for 3 days. Then, we mixed 80% of this enriched flour with 20% of fresh wheat flour to ensure that nutrient depletion of the enriched flour would not lead to starvation in the recipients. We placed single recipient pupae collected from our stock population in 1.5 mL tubes containing either 0.5 g of fresh wheat flour (control) or flour enriched with larval or adult feces. Every week, we sampled and stored individuals (n = 10 females/week/treatment) for further processing (Fig S4B).

#### Experiment 8

Finally, we tested whether the amount of larval fecal inoculum would alter recipient beetle microbiomes. We generated larval-enriched wheat flour as described above (Experiment 7), and then prepared a series of flour mixes with different proportions of enriched flour (0%, 25%, 50%, 75%, 100%), adding fresh wheat flour as required. We placed single pupae in each treatment as described above for 3 weeks, and sampled at the end point (i.e., 2-week-old adults) for microbiome analysis (Fig S4C).

### Measuring the impact of quinones on bacteria and the beetle microbiome

#### Experiment 9

Adult flour beetles secrete antimicrobial quinones from specialized stink glands (18,19), and we suspected that this may explain why adult presence reduced flour bacterial loads and increased individual level microbiome variation (see Results). To test whether beetle-associated bacteria are differentially sensitive to quinones, we first isolated bacteria from beetle life stages and used flour, using multiple solid growth media (Table S1). To prepare glycerol stocks, we scraped colonies from the plate, resuspended them in 800 µL of autoclaved 60% glycerol solution, and stored at −80°C. We extracted genomic DNA for each isolate, and identified it by Sanger sequencing the 16S rRNA gene (Table S1). To prepare quinone extracts, we took 100 6-week-old adult beetles from the stock population and placed them in an empty Petri plate (90 mm diameter). We stimulated the release of quinones from stink glands using a cold shock, placing the plate on ice until beetles were immobile. We removed the plate from the ice, allowed beetles to recover, and repeated the process for a total of 30 mins. We removed the adults and collected the released quinones by adding 1 mL of hexane to the plate. After mixing with a pipette, we transferred the quinone extract to a 1.5 mL tube. We revived bacterial stocks by streaking on Luria-Bertani (LB) agar (Difco) and M-Enterococcus agar (HiMedia) plates, incubated at 28°C for 24 hrs, picked up multiple colonies, suspended in 0.9% autoclaved saline and normalized all cultures to an OD_600_ of 0.1 using sterile saline. We spread the culture on 1.5% LB agar plates using sterile swabs to make a bacterial lawn, and dried the plates inside a laminar hood for a few minutes. We placed 4 sterile filter paper discs (6 mm diameter) onto the prepared bacterial lawn plates, spaced evenly apart. We added 20 µL of the prepared quinone extract onto 3 of the discs and the solvent (hexane) as our negative control for the 4^th^ disc (1 Petriplate per isolate, with 3 technical replicates of the test and 1 control). We incubated plates at 28°C for 14-16 hrs (∼40 hrs for *Enterococcus* isolates because of slow growth), and measured the zone of inhibition around each disc as a measure of sensitivity to quinones.

#### Experiment 10

We tested whether focal bacterial strains are also sensitive to quinones secreted in wheat flour. We took fresh wheat flour and added feces (which are rich in the beetle-associated *Izhakiella* strain that we are unable to culture), or 0.3% lyophilized bacterial cells (*Acinetobacter radioresistens* or *Enterococcus faecium*) (20), either with or without quinone extract. The treatments without quinones allowed us to measure bacterial death in the absence of quinones. We prepared quinone extract (as described for Experiment 9) and added 50 µL extract to 0.5 g of flour in 1.5 ml tubes with mixing, every 4^th^ day. We periodically stored a subset of tubes at −80°C to measure the bacterial death rate (n = 8 replicate tubes/treatment/time point) (Fig S5A).

Next, we tested whether quinones perturb microbial load in the flour in the presence of beetles, which the flour bacteria could colonize. We prepared flour enriched with larval or adult feces, as described for Experiment 7. In another treatment, we added both larval feces and quinone discs. However, here the quinone preparation differed slightly: we placed 100 beetles directly in hexane at −80°C (without a cold shock), and added quinone extract to the flour only at the beginning of the experiment. We placed single pupae isolated from the stock population in a 1.5 mL tube containing 0.5 g of the appropriate flour for each treatment. We sampled a subset of pupae every week for 6 weeks, storing 10 females/week/treatment at −80°C for further processing (Fig S4B).

#### Experiment 11

To test the impact of quinones on the microbiome, we used RNA-mediated interference (RNAi, using double-stranded RNA, dsRNA) to knock down expression of the gene *GT39*, which is involved in quinone synthesis (21). As negative controls, we included beetles that were left untreated, injected with TE buffer (10mM Tris, 1mM EDTA buffer to control for effects of injection), or with GFP (green fluorescent protein) dsRNA (which should not influence quinone synthesis). To synthesize *GT39* dsRNA, we extracted DNA from female beetles (as described below), and amplified the gene of interest using primers with the T7-promoter, described in (21).

GT39_Forward-5’-TAATACGACTCACTATAGGG-GGAGGTCACCCAGAACAACT-3’; GT39_Reverse-5’-TAATACGACTCACTATAGGG-TGACATCCCTTGGCACATATTC-3’; To obtain the dsRNA for GFP, we used a strain of *Escherichia coli* MG1655 carrying GFP to get template DNA, and used the following primers that we designed.

GFP_Forward-5’-TAATACGACTCACTATAGGG-GCAGCCGGATCCTTTGTATAG-3’; GFP_Reverse-5’-TAATACGACTCACTATAGGG-TGCTGAAGTCAAGTTTGAAGGTG-3’. We used 500 ng of the amplified DNA as a template for in-vitro transcription in a 20 µL reaction using the Thermo MEGAscript RNAi Kit. We followed the kit instructions, with overnight incubation at 37°C followed by elution of the dsRNA in TE buffer. We injected 0.2 µL of the dsRNA solution (∼200 ng/pupae) in early-stage (∼2 day old) pupae using a microinjector. We handled control pupae in an identical fashion, except untreated controls which were not injected. We isolated pupae in 1.5 mL tubes containing 0.5 g of fresh wheat flour for one week, during which time the pupae eclosed into adults. We grouped these adults into 10 pairs (10 individuals per sex) in 3 g of flour in a small plastic box (35 mm diameter) for 3 weeks (i.e., until we obtained ∼4 week old adults) (n = 7 replicate boxes/treatment). One week after grouping, we sampled and dissected a few females to check the stink gland phenotype (n = 2 females from each of 3 replicate boxes per treatment). At the final time point, we stored all remaining beetles until further processing (Fig S5B).

### Measuring the impact of periodic flour replacement on the beetle microbiome

#### Experiment 12

Our results suggested that fresh flour has very little bacterial inoculum, that fecal inoculum from larvae is critical for the colonization of the beetle gut, and that adult-secreted quinones selectively suppress the growth of specific bacteria. This led us to predict that in our stock populations, substantial ecological drift may be generated due to a combination of these factors and our stock maintenance protocol, which involves periodic replacement of the flour. To test this, we sampled and stored flour (0.5 g) and beetles (n = 10 for each available life stage) every 5 days from our main stock population. We sampled and analysed microbiomes across two successive discrete-generation cycles. We also similarly sampled an independent experimental population that was maintained under continuous generation cycles, in which the number of larvae does not fluctuate as much as under discrete generations.

### Measuring the effect of microbiome on beetle fitness

#### Experiment 13a

Our prior work ((17) suggested that the flour microbiome is beneficial for beetles, increasing fitness parameters such as female fecundity. We repeated this experiment several times, following the protocol described in Agarwal et al to disrupt the flour microbiome using UV irradiation (exposing thin layers of flour to UV in a laminar hood for 2 hours) or addition of antibiotics. Briefly, we placed mated females from the stock population in ∼0.5-0.7 g of double sifted fresh or treated flour and counted the number of eggs laid by each female after 48 hrs as a measure of fitness. Note that Agarwal et al used single antibiotics at a time (0.005% w/w), but we used a combination of three antibiotics (0.05% w/w each of kanamycin, streptomycin, and tetracycline).

#### Experiment 13b

Next, we tested whether the addition of larval feces could increase beetle fitness (Fig S6A). We collected eggs from a stock population, and reared them in groups of 500 in 500 g of either fresh wheat flour or fresh flour enriched with larval feces (50% used flour, generated as described for Experiment 7 above) to adulthood. We measured the fecundity of 2-week old females provided either 3g fresh flour or flour enriched with larval feces (50% fresh flour), as described above.

### DNA extraction and sequencing for flour and beetle samples

We extracted DNA from stored (–80°C) or fresh beetles (larvae, pupae, or adults) using the Qiagen DNeasy Blood and Tissue kit, with the following modifications. We first surface-sterilized each sample by dipping it in 70% ethanol, and then washed it with sterile water. For adult beetles, we removed the wings (elytra) and legs. The beetles were again surface sterilized, washed, and kept in individual 1.5 mL tubes. We flash froze each beetle in liquid nitrogen and crushed it using a micro-pestle. Next, we added 180 µl of lysis buffer and 20 µl Proteinase K solution from the kit, vortexed the sample, and kept it at 56°C for ∼18 hours. Then, we followed the rest of the kit protocol, eluting DNA in 30 µl of nuclease-free water.

To obtain the wheat flour microbiome, we modified the protocol to increase the yield of bacterial DNA and minimize plant (flour) DNA. We took 100 mg of wheat flour, added 300 µL of Redford buffer (1 M tris HCl; 0.5 M EDTA; 1.24% Triton), and placed the tube horizontally with periodic gentle shaking on a vortex for 2 hrs, as described in (22). We pelleted the flour debris by centrifuging at 10,000 rpm for 5 mins, and used the supernatant for further processing as described above.

We quantified the DNA concentration in each sample using the Qubit dsDNA HS assay kit. We amplified the V3-V4 hypervariable region of the 16S rRNA gene using a modified Illumina 10N primer to facilitate detection of rare taxa (23). We followed the standard Illumina 16S metagenome amplification protocol, with the following modification. For wheat flour and larval samples, which have high chloroplast and mitochondrial contamination, we added 25 and 15 µM PNAs, respectively (peptide nucleic acids, to specifically block chloroplast and mitochondrial DNA amplification; (24). We prepared libraries from the amplified product as per standard Illumina protocols and sequenced on the Illumina Miseq platform (300 x 2 paired-end reads). In each MiSeq flow cell, we also included one negative control, where we did not add any DNA template, but carried out all amplification steps.

### Measuring total bacterial load using quantitative PCR

We measured absolute bacterial load in the same beetle and flour samples used for microbiome analysis (n = 6-10 for beetles and 2-4 for wheat flour). We used 1 µL of extracted DNA solution from each sample (described above) and obtained 200 bp amplicons for quantification, using primers designed to amplify only bacterial 16S rRNA (forward: 5’-CGGTAATACGGAGGGTGCAA-3’; reverse: 5’-TCCTCCAGATCTCTACGCAT −3’), using the qPCR protocol described in (22). For beetle samples, we normalized the amount of bacterial DNA to the amount of beetle DNA, using primers for the host gene *rps3* (not found in bacteria or plants) (forward 5’-CAGGGGTCTGTGTGCCATTG-3’; reverse 5’-TCACACCCTGCAAAATAACCACT-3’). Similarly, we normalized flour samples to the actin gene *ACT2* (forward: 5’-CAAATCATGTTTGAGACCTTCAATG-3’; reverse 5’-ACCAGAATCCAACACGATACCTG-3’) (25) that represents the amount of plant DNA. In this manner, we effectively estimated the total bacterial load per gram host tissue (or flour), as 2^-ΔCt^, where ΔCt = Ct(16S rRNA) – Ct(rps3 or actin).

### Data analysis

We used R for all statistical analysis (26). To determine the microbiome composition from the amplicon sequencing output from MiSeq, we used the DADA2 pipeline (27) to generate a taxonomic table using 100% identity with the Silva reference database (training set v138.1) (ref). We first filtered out chloroplast and mitochondrial reads, obtaining an average of 30,000 reads (range 20,000-60,000) in beetle samples, and 5,000 reads (range 5,000-15,000) in flour samples. We obtained less than 100 reads on average in our negative controls, suggesting no significant contribution of contaminants from reagents. We used a subset of samples to conduct a rarefaction analysis, finding that ∼10,000 bacterial reads for the beetle samples, and ∼5,000 bacterial reads for the flour samples, led to saturation of community richness respectively (Fig S7). For samples with fewer reads than these thresholds, we repeated the sequencing. Samples that repeatedly failed to generate sufficient reads were excluded from further analysis.

We used the R packages ‘phyloseq’ and ‘vegan’ to estimate Shannon diversity in all samples (28,29). To quantify individual variation in microbiome composition, we used the package ‘NST’ (30) to calculate the taxonomic beta diversity (β_rc-_ Raup-Crick Dissimilarity) (31). We carried out PERMANOVA using the ‘adonis2’ function in the package ‘vegan’ in R to quantify the impact of various factors on the beetle microbiome. Finally, we used the R package ‘ggplot2’ for visualization.

## RESULTS

### Beetle microbiomes vary across time, replicate populations, generations, and individuals

We first analyzed the microbiomes of female beetles sampled from the same outbred stock population across years (Experiments 1a and 1b, Table 1). The bacterial community composition varied significantly across time (Fig 1A; PERMANOVA, timepoint effect (month/year combination): R^2^ = 0.25, P = 0.0009; year effect: R^2^ = 0.13, P = 0.0009), as did bacterial richness and diversity (Fig S8; Kruskal-Wallis tests, P_diversity_ = 0.01; P_richness_ = 6.5 e-05). We observed similar patterns across three sampled time points for males, indicating that underlying processes were not sex-specific (Fig S9). Next, we tested whether three independently maintained stock populations initiated from the same founders also acquired divergent microbiomes (Experiment 1c, Table 1). In February 2016, the populations had indeed diverged, although we could not identify the dominant bacterial taxon (Fig 1B; PERMANOVA, lineage effect: R^2^ = 0.13, P = 0.003). In March 2022, the three populations appeared to harbour more similar microbiomes dominated by *Izhakiella*, though the community composition still differed significantly (Fig 1B; lineage effect: R^2^ = 0.10, P = 0.004). Males at this time point showed similar patterns (PERMANOVA, lineage effect: R^2^ = 0.11, P = 0.0009, Fig S9). Thus, in addition to temporal changes observed in a given population over several years, the microbiomes of independently maintained stock populations also diverged significantly from each other.

**Fig 1:**
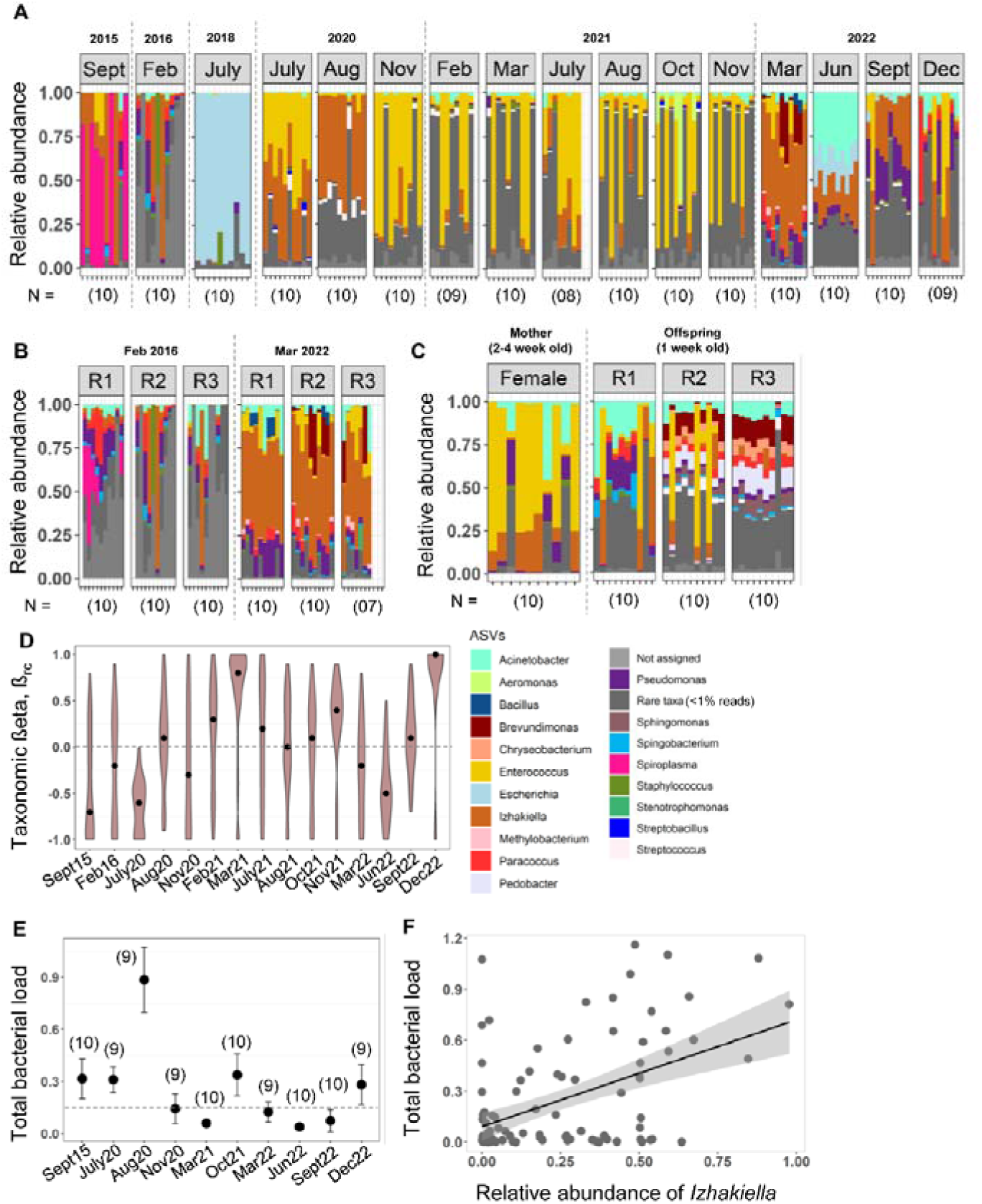
Beetle microbiome and bacterial load varies across time, individuals, populations, and generations. Results from Experiments 1 and 2 are shown here. (A–C) The relative abundance of bacterial ASVs (identified to genus level) for each female beetle sampled from a population as a function of (A) time (B) replicate populations (three different lineages) (C) replicate populations from the same stock population. (D) Taxonomic beta diversity (β_rc_) of bacterial communities across time, for samples shown in panel A. Positive values indicate that samples are more dissimilar; negative values indicate that samples are taxonomically more similar to each other. (E) Mean total bacterial load per beetle (normalized to total amount of host DNA per sample) for a subset of samples shown in panel A. The dotted grey line represents the overall average bacterial load across all samples, excluding August 2020 samples. Error bars represent standard error. (F) Correlation between the mean relative abundance of *Izhakiella* and the mean total bacterial load per population, for samples shown in panel E. The trendline shown is a linear model fit with R^2^=0.22. Sample size (number of female beetles) is indicated in parentheses.

**Table 1:** Summary of experiments conducted, and corresponding figures.

| Experiment | Description | Figures |
| --- | --- | --- |
| 1a,b | Quantify temporal variation in microbiomes: compare microbiomes of adults from a single stock population stored at -80° C from 2015–2022 | Fig 1A,D-F, Fig S8-S11 |
| 1c | Quantify microbiome variation across independently evolving lineages: compare microbiomes of adults from independent stock population boxes founded by same parental stock, sampled at the same time | Fig 1B |
| 2 | Quantify microbiome variation across replicate population boxes: compare microbiomes of adult offspring across 3 replicate boxes set up using the same stock parents | Fig 1C, Fig 4D-F |
| 3 | Determine microbiomes as a function of external factors such as experimenter identity, air sterility, and fresh vs. heat-treated wheat flour | Fig S12 |
| 4a | Test for maternal microbiome transmission, comparing microbiomes of females and eggs | Fig S13A |
| 4b | Test for maternal microbiome transmission, analysing microbiomes of dissected female eggs and ovaries | Fig S13C |
| 4c | Test for consistent maternal microbiome transmission, | Fig S13D |
|  | comparing microbiomes of eggs dissected from ageing females |  |
| 5 | Quantify effect of population density: comparing microbiomes of adult offspring in parental populations housed at two different densities | Fig 2A-C, Fig S14 |
| 6 | Quantify fecal microbiomes of larvae and adults across development | Fig 3A-B, Fig S15A-B |
| 7 | Quantify effect of fecal microbial inoculum on adult microbiomes, comparing adults provided with fresh flour, flour enriched with larval vs. adult feces, or flour enriched with larval feces and added quinones | Fig 3C-D, Fig S145C |
| 8 | Quantify effect of amount of larval feces on beetle microbiome: comparing microbiomes of adults fed with different proportions of larval feces in fresh flour | Fig 3E-F, Fig S15D-E |
| 9 | Test the sensitivity of beetle-associated bacterial strains to adult-secreted antimicrobial quinones | Fig 4A |
| 10 | Test the effect of quinones on bacterial load in flour: comparing temporal change in flour microbial loads, after inoculating with lyophilized focal bacteria or larval or adult feces, with vs. without quinones | Fig 4B, Fig S16A |
| 11 | Test the effect of adult quinones on beetle microbiomes: comparing microbiomes of beetles with vs. without RNAi-mediated knockdown of quinone pathway | Fig 4C-D, Fig S16C |
| 12 | Test the effect of flour replacement on the flour microbiome: comparing flour microbiomes across two cycles (generations) for stock populations maintained under discrete or continuous generations. | Fig 5A-C, Fig S17A-D |
| 13a | Test the effect of depleting the microbiome on beetle fitness. | Fig 6 |
| 13b | Test the effect of enriching the flour microbiome with larval feces on beetle fitness, after rearing them (from eggs) on enriched flour. | Fig S6 |

Next, we tested for shorter-term variation in microbiome composition (Experiment 2). For three replicate “offspring” boxes set up from the same set of stock-derived parents, we again observed significant variation in microbiomes (PERMANOVA, replicate effect, females: R^2^ = 0.23, P=0.0009; males: R^2^ = 0.22, P=0.0009). Thus, beetle microbiomes varied dramatically across time and across independent lineages maintained under the same laboratory conditions. Despite this variation, however, a few bacterial genera such as *Izhakiella*, *Enterococcus*, *Pseudomonas*, and *Acinetobacter* often dominated the community (Fig 1A), and were identified as members of the core beetle microbiome (Fig S10). For instance, *Izhakiella* and *Enterococcus* were consistently present in ∼80% of beetle samples with a moderate relative abundance of 10%, whereas *Pseudomonas* and *Acinetobacter* were found in ∼50% of the samples at a low relative abundance of 1% (Fig S10).

We then analysed the degree of individual-level variation in microbiome composition, which also appeared to be high (Fig 1A). Using taxonomic beta diversity (ß_rc_) as a metric of microbiome diversity, we found that the degree of individual variation in a population also changed across time (Fig 1D; Kruskal-Wallis test, time effect: P < 2.2e-16). For instance, females had highly similar microbiomes in September 2015, July 2020, and June 2022 (median ß_rc_ approaching −1), whereas females sampled from the same stock population had highly dissimilar microbiomes in March 2021 and December 2022 (median ß_rc_ approaching +1) (Fig 1D). We also tested whether individual variation in microbiome composition was associated with total bacterial load, measured using qPCR (normalized with a beetle-specific primer to obtain an estimate of total bacterial DNA per unit host DNA for each sample). The bacterial load per beetle varied significantly over time (ANOVA, load ∼ time: P = 4.64e-06 Fig 1E), but was not correlated with ß_rc_ (Spearman’s rho = −0.14, P = 0.16). Both results were robust to removing a potential outlier data point from August 2020 (ANOVA, load ∼ time: P = 0.012; Spearman’s rho = −0.15, P = 0.17). In contrast, the total bacterial load was positively correlated with the relative abundance of the genus *Izhakiella*, suggesting that this genus may colonize beetles consistently and well (Fig 1F; Spearman’s rho =0.34, P = 0.0002). We did not observe such correlations for the relative abundance of *Enterococcus* or for rare taxa that sometimes had a relatively high combined relative abundance (e.g., Fig 1A, March 2022 samples) (Fig S11).

Together, these results showed that the beetle microbiome composition and load vary a lot across individuals, generations, and populations; but dominant taxa recur, including the genus *Izhakiella,* which appears to be a strong colonizer. Next, we considered factors that could drive this variation across individuals and over time. The flour beetle system is a relatively closed environment, where the flour serves as both habitat and food for all life stages. All individuals consume flour and deposit fecal material in the flour (except pupae, which do not feed), such that microbes continuously circulate through beetles and flour, enabling dispersal and exchange across host individuals. Hence, there are broadly three sources of bacterial inocula in this system: the environment, flour, and beetles. We considered each of these in turn.

### Sources of bacterial inoculum: external environmental factors and fresh flour

We first tested whether variation in external sources of bacterial inocula such as air, experimenter, and fresh flour could contribute to the variation in beetle microbiomes. We set up independent experiments to test these factors, with the following treatments: exposure to normal vs. sterile air; experimenter 1 vs experimenter 2; and fresh flour vs. heat-treated fresh flour (Experiment 3). In each case, we found relatively weak effects of the treatment. Air sterility had no significant effect, experimenter identity explained ∼15% of the variation in bacterial community composition, and flour treatment explained ∼13% of the variation (Fig S12A) in adult female microbiomes. All treatments had a significant effect on adult male microbiomes but only ∼13%-16% of variation was explained by these factors (Fig S12B). All factors also influenced the degree of individual variation (Fig S12C; Wilcoxon tests, P_air sterility_ = 0.0001; P_experimenter_ = 0.003; P_flour_ = 0.0004). For instance, experimenter 1 generated individuals with more similar microbiomes compared to experimenter 2, and individuals reared on dry heated wheat flour had highly dissimilar microbiomes compared to those fed fresh wheat flour. Although air sterility and flour affected the relative abundance of *Izhakiella* (Pearson chi-square test, P_air sterility_ = 0.005546, P_flour_ = 4.531e-09; Fig S12A), none of the factors influenced total bacterial load (Fig S12D; ANOVA, load ∼ factor, P_experimenter_= 0.3; P_air_ = 0.6; P_flour_ = 0.3), which tended to be lower than that observed in our temporal sampling experiment (compare Fig S12D with Fig 1E). The experimenter identity effect was potentially confounded by an unexpected difference in population density between the boxes handled by the two experimenters (e.g., experimenter 1 had ∼700 larvae whereas experimenter 2 had ∼1200 larvae on the 18^th^ day). This density effect was specifically tested later. Together, our results suggested that the tested external environmental factors are unlikely to explain the observed large variation in beetle microbiomes, though it is possible that these factors may be important in combination and/or over several generations.

### Sources of bacterial inoculum: host-associated factors

Next, we investigated the role of possible host-associated sources of bacterial inoculum on the beetle microbiome, which may include maternal transmission through the egg, and horizontal transmission via larval or adult beetle feces. In a preliminary experiment, we observed that microbiomes of groups of freshly laid eggs were different from those of their mothers (Experiment 4a, PERMANOVA, R^2^ = 0.09, P = 0.002, Fig S13A) with lower abundance of *Izhakiella*, in contrast to expectations under maternal transmission. For eggs and ovaries dissected from females before oviposition (Experiment 4b, Fig S13B), we again observed very low relative abundance of *Izhakiella* (∼3%, Fig S13C). Finally, we found that the egg microbiome varied significantly with maternal age (Experiment 4c; PERMANOVA, age, R^2^ = 0.35, P = 0.014; Fig S13D), indicating unstable egg microbiomes. Together, these results suggest a lack of substantial and consistent bacterial transmission from mother to offspring via eggs.

Therefore, we turned to the possibility that feces could serve as an important source of bacteria that colonize beetle guts. If true, higher population densities should reduce the stochasticity in colonization by increasing fecal inoculum in the flour. Recall that our previous results from Experiment 3 (different experimenters happened to generate populations with distinct densities, which also had different microbiomes) are consistent with this prediction. Hence, we set up a new experiment where a single experimenter set up beetle populations at either low or high adult density. Here, we observed that compared to low density populations, individuals reared in high density populations had similar total bacterial loads (ANOVA, load ∼ density, P = 0.06, Fig 2A) but higher relative abundance of *Izhakiella* (average 47% vs. 18% in high vs. low density respectively), lower individual variation (Wilcoxon test, P = 3.10e-14), and distinct microbiomes (Experiment 5, PERMANOVA, density effect combining all replicates, R^2^ = 0.06, P = 0.0009; Fig 2B-C, Fig S14). Replicate populations within each treatment also varied significantly (PERMANOVA, replicate effect: low density, R^2^ = 0.17, P = 0.0009; high density, R^2^ = 0.13, P = 0.0009), potenitially reducing the effect of population density. Nonetheless, these results pointed to fecal inoculum as an important factor affecting microbiome variability.

**Fig 2:**
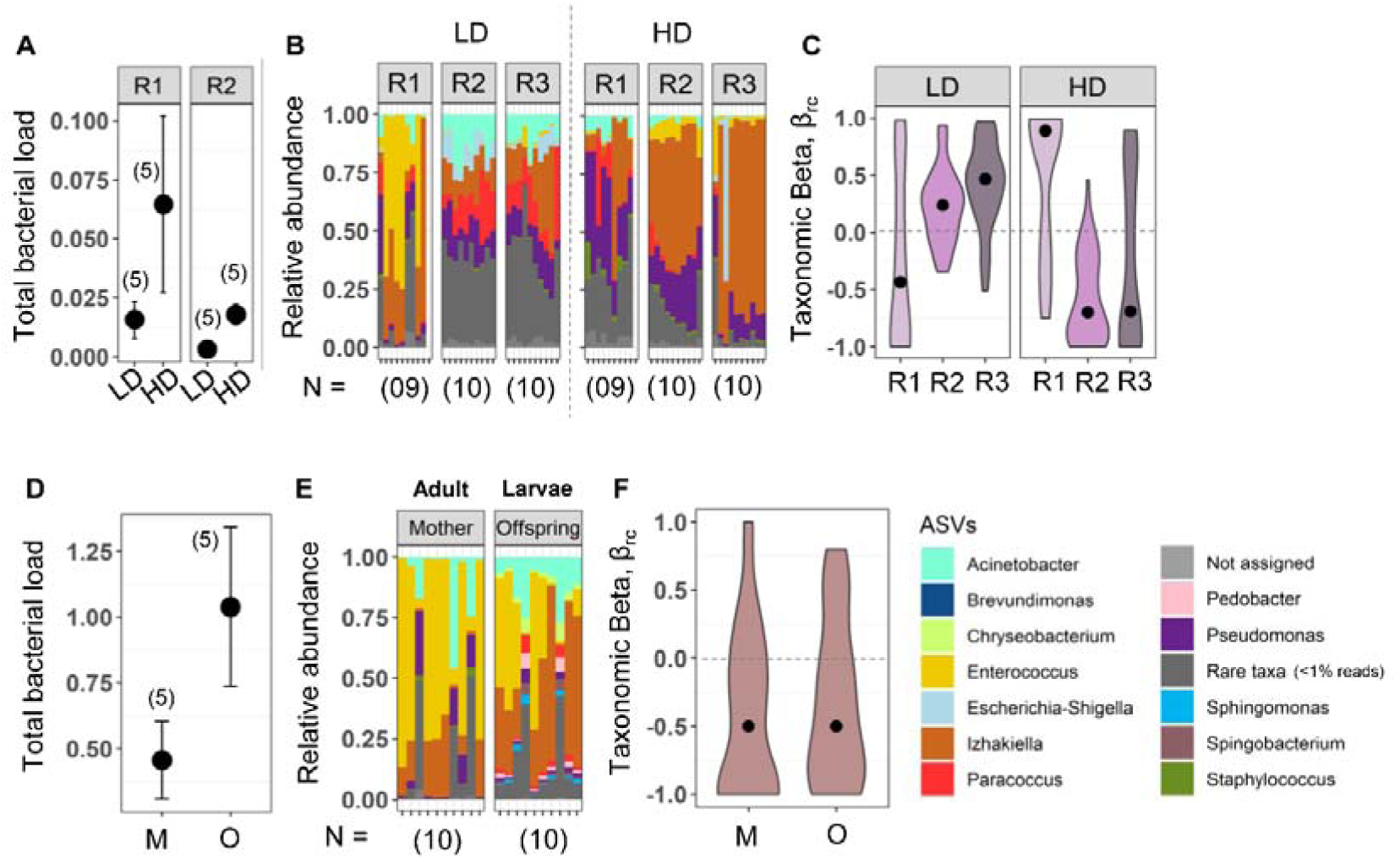
Feces are a major source of beetle microbiome inoculum. (A–C) Results of Experiment 5 determining the bacterial community characteristics of the offspring of 2-week-old females ovipositing at different adult densities (LD = low density; HD = high density; R1, R2, R3 = replicates). (A) Mean total bacterial load (±se) (B) Bacterial community composition (C) Taxonomic beta diversity (β_rc_). (D–F) Results from Experiment 2 comparing bacterial communities of females and their 14-day old larval offspring. (D) Mean total bacterial load (±se). (E–F) Bacterial community composition (E) and Taxonomic beta diversity (β_rc_) (F) of adult females (M = mother) and 2 week old larvae (O = offspring). Sample sizes are indicated in parentheses.

In the above experiment, we focused on adult fecal material, but larvae are also a potentially important source of fecal microbial inoculum. On one hand, the microbiomes of larvae and adults could be similar since all life stages consume the same flour, which includes fecal material from all individuals in the population. On the other hand, given the distinct feeding patterns and physiology of larvae and adults, distinct microbiomes could be associated with each life stage. Indeed, we found significant differences in both the microbiome composition and load of larvae vs. adults sampled from the same populations (Experiment 3, PERMANOVA comparing larvae vs. parents; R^2^ = 0.21, P = 0.0009; Fig 2D–F). Larvae had significantly higher abundance of *Izhakiella* (two experimental blocks: 30% vs. 14% in larvae and adults respectively, Chi-square test, P = 0.01; and 53% vs. 6%, P = 9.42e-10). Given that the larval stage is the primary growing stage and larvae consume a large amount of flour to support the rapid increase in body size, we speculated that larval feces may serve as a major source of bacterial inoculum in the flour.

We tested this by sampling feces of larvae vs. adults across development. The larval fecal microbiome composition was very stable and was primarily composed of *Izhakiella* (∼80%) with the highest bacterial load at days 9-10, which is approximately halfway through the total larval period of ∼20 days (Experiment 6, Fig 3A, Fig S15A). In contrast, the adult fecal microbiome was highly variable until 2 weeks post-eclosion, and then stabilized with a large increase in total bacterial load (Fig 3B, Fig S15B). Overall, the bacterial load in larval feces was higher than in adult feces (ANOVA, load ∼ age x stage, P_age_ = 3.52e-6; P_stage_ = 0.0007; P_age x stage_ = 0.0007). Given that most of our stored adult beetle samples (analysed in the first section of the Results) were ∼2 weeks old, these results may help explain why we observed high variability in beetles sampled across time (Fig 1A). Further, exposing pupae to larval feces led to higher bacterial loads, more stable microbiomes, and lower individual variation in the eclosed adults, compared to pupae exposed to flour enriched with adult feces (Experiment 7, Fig 3C-D, statistics reported in the figure legend, Fig S15C). The stabilizing effect of larval feces also increased with the proportion of fecal inoculum provided in the flour (Experiment 8, Fig 3F-G, statistics reported in the figure legend, Fig S14D-E). Together, these results support a strong role of fecal-oral transmission in microbiome assembly and stability, with larval feces as the main source of inoculum for the beetle microbiome.

**Fig 3:**
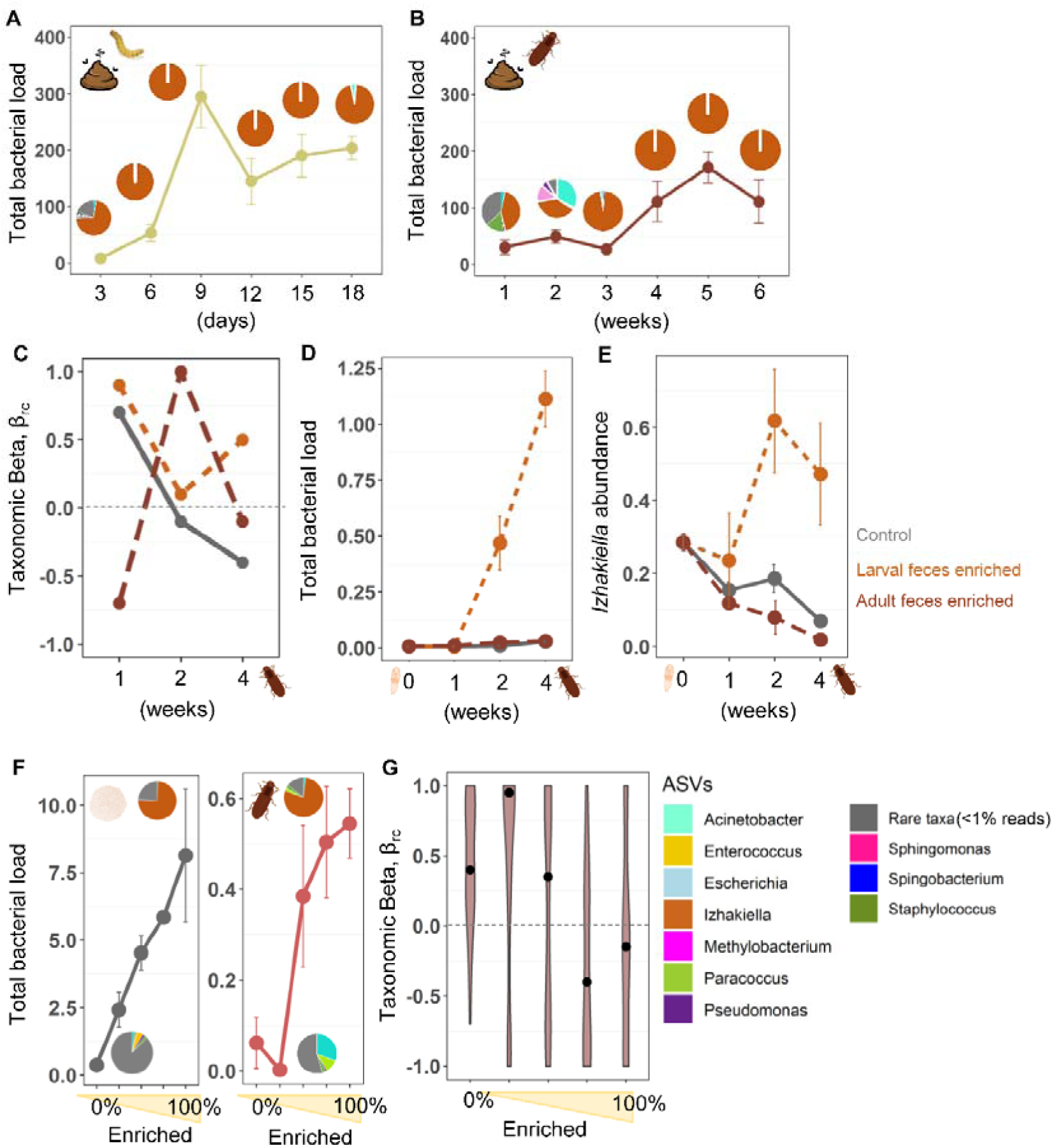
Source and amount of host fecal inoculum in flour alter the beetle microbiome. (A–B) Results of Experiment 6 to determine the time course of mean total bacterial load (±se) and average relative abundance of bacterial taxa (shown as pie charts) in the feces of (A) larvae and (B) adults (n = 3 replicate tubes, each containing 30 larvae or adults) (see Fig S13A-B for microbiome composition). (C–D) Results of Experiment 7 to determine microbiome establishment in pupae exposed to flour enriched with larval or adult feces, or nothing (control), as a function of the age of eclosed adult females. (C) Median taxonomic beta diversity (β_rc_) (see Fig S13C for microbiome composition) (ANOVA, P_treat_ = 0.79, P_time_ = 0.009, P_treat X time_ = 1.19e-06) (D) Mean total bacterial load and (E) relative abundance of *Izhakiella* (n = 6 females/treatment/age) (ANOVA, P_treat_ = 0.006, P_time_ = 0.003, P_treat X time_ = 0.0002). (F–G) Results of Experiment 8 to test the effect of amount of fecal inoculum on beetle microbiome. (F) Mean total bacterial load in the flour (n = 2 technical replicates/treatment) (ANOVA, P_treat_ = 0.03) and in beetles (n = 4 females/treatment) (ANOVA, P_treat_ = 0.003). Pie charts show the average bacterial community composition in 25% and 100% enriched flour (see Fig S13D for the full dataset). (G) Taxonomic beta diversity (β_rc_) (see Fig S13E for full dataset) (Kruskal-Wallis tests, P_diversity_ = 9.586e-06). Error bars represent standard error.

### The role of adult-secreted quionones

The results above indicated that fecal inoculum is important, and that it is life-stage specific. However, beyond the effect of fecal volume and turnover in the gut (which is expected to be higher for larvae), we suspected that adults also have an inhibitory effect on bacterial load in the flour, given prior work showing antimicrobial effects of quinone compounds secreted from adult stink glands in the flour (18,19). The shift in adult microbiomes between 2-3 weeks of age coincides with the stink glands becoming fully functional (32,33). Further, the effect of adult density observed above (higher density reduces individual microbiome variation) could be mediated by density-dependent upregulation of quinones (34). Hence, we tested whether bacteria in the beetle microbiome were sensitive to adult-secreted quinones (Experiment 9).

We found differential sensitivity of beetle-associated culturable bacteria to stink gland extracts from 6-week old females; the solvent negative control did not inhibit growth (Fig 4A, ANOVA, zone of inhibition ∼ isolates, P = 0.001). Different isolates of the same genus also had distinct sensitivity (e.g., *Enterococcus* and *Acinetobacter*), demonstrating large functional variation even in closely related bacteria. Notably, opportunistic pathogens such as *Serratia* were among the most resistant to quinones. Unfortunately, we were unable to culture *Izhakiella* from beetles or larvae; hence we used *I. capsodis* that was previously isolated from mirid bugs (35). This species was very sensitive to quinones, but given the variation observed among other congeners, it is unclear whether our beetle-associated *Izhakiella* strain(s) have similar quinone resistance as *I. capsodis*.

**Fig 4:**
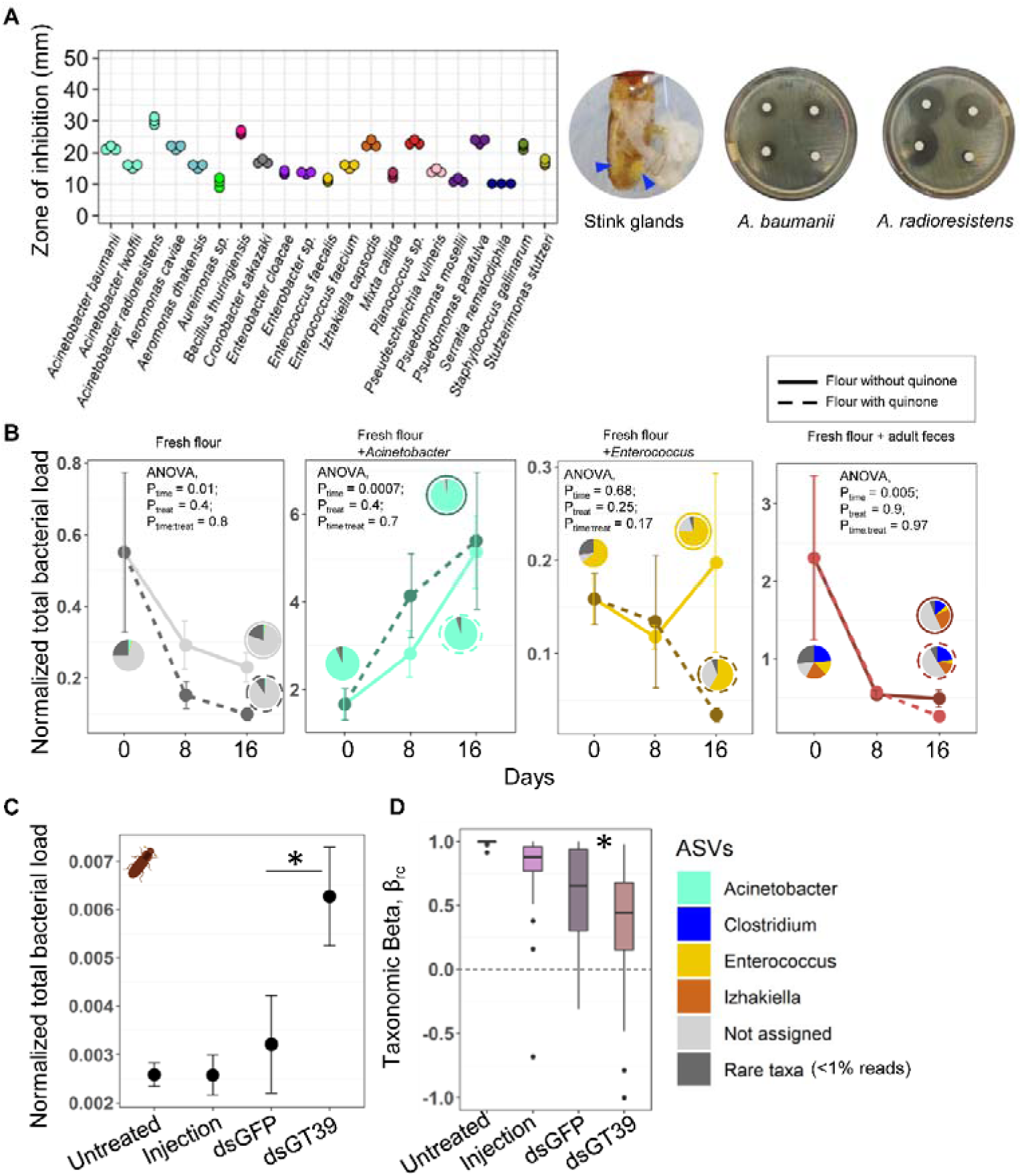
Effect of quinones on the adult microbiome. (A) Results of Experiment 9 to measure the antimicrobial effect of quinone extracts on several bacterial strains isolated from beetles (n = 3 technical replicates/strain). A larger zone of inhibition indicates greater sensitivity of the strain to quinones. The images show a dissected beetle with paired abdominal stink glands (blue arrows), and examples of agar plates with bacterial growth inhibition due to quinones added to filter paper discs (the bottom right disc in each plate serves as a solvent-only negative control). (B) Results of Experiment 10 showing the effect of quinones on mean (±se) total bacterial load in the flour (n = 8 technical replicates/treatment/timepoint). Results of ANOVA are shown for each panel. Pie charts show the average bacterial community composition; outer circles indicate treatment (solid line: without quinone; dotted line: with quinones) (see Table S2 and Fig S14A for data for each replicate). (C–D) Results of Experiment 11, testing the effect of suppressing quinones using RNAi for genes GT39 (quinone production) and GFP (negative control); with untreated beetles and injection with buffer serving as additional controls. (C) Mean (±se) total bacterial load in beetles (n = 4 females/treatment) (t test). (D) Taxonomic beta diversity (β_rc_) of experimental beetles (see Fig S14C for microbiomes of individual females). Asterisks indicate significant differences between dsGT39 and dsGFP beetles (Wilcox test).

We next tested whether quinones influence bacterial persistence in the flour, in the absence of live adult beetles (or larvae) (Experiment 10). We added large quantities of lyophilized bacteria or adult fecal material to flour, and monitored bacterial loads over time, with or without added quinone extracts. Quinones had differential effects on the load of different bacteria in flour (Fig 4B, Table S2). The total bacterial load in fresh flour as well as flour enriched with adult feces generally declined over time (likely due to dessication in the dry flour environment), and the decline was exacerbated on adding quinones (Fig 4B, Fig S16A). However, lyophilized *Enterococcus* survived for at least ∼2 weeks in flour, and declined only when we added quinones. Surprisingly, the load of *Acinetobacter* increased significantly over time (Fig 4B, Fig S16A). We were unable to identify the cause of this counterintuitive bacterial growth, which is not expected to happen in dry flour. Nonetheless, the decline in bacterial load due to quinones was robust to a different method of quinone extraction (Fig S16B). Thus, adult secreted quinones could play an important role in modulating bacterial loads in flour, potentially influencing microbiome assembly and dynamics.

To test this hypothesis, we suppressed quinone production in adult beetles via RNA-mediated interference of expression of the gene *GT39*, which is involved in quinone synthesis (21) (Experiment 11). The dsGT39 treated beetles had missing or underdeveloped stink glands, confirming that the RNAi was effective (Fig S5C). Compared to controls, the bacterial load in these beetles was significantly higher, as expected (Fig 4C, t test, dsGFP vs. dsGT39: P = 0.014). The quinone knockdown beetles also had significantly lower individual variation in microbiomes (Fig 4D, dsGFP vs. dsGT39, Wilcoxon test, P = 0.02). However, the microbiome composition of beetles was similar in both cases (PERMANOVA, dsGFP vs. dsGT39: R^2^ = 0.05, P = 0.4), and we did not observe high *Izhakiella* abundance in any of the beetles (Fig S16C). Thus, the effect of quinone production was weaker than we expected, perhaps due to the lack of initial larval fecal inoculum in the fresh flour supplied to pupae in this experiment. Nevertheless, these results demonstrate an important role for adult-secreted quinones in regulating flour beetle microbiomes.

### Population maintenance in the laboratory affects flour microbiomes and bacterial load

Our experiments thus far highlighted two important factors driving beetle microbiome assembly and stability: larval fecal inoculum, which increases bacterial loads in the flour, and adult-secreted quinones, which suppress it. However, the periodic replacement and replenishment of flour — a critical part of flour beetle population maintenance — could potentially dilute the effect of these two factors, and therefore also influence microbiomes. As individuals grow and fecal matter accumulates, the flour quality declines, flour can get moldy, and populations become more susceptible to infection and nutrient deficiencies. Therefore, periodically replacing conditioned flour with fresh flour is essential for maintaining healthy laboratory populations of flour beetles.

In our maintenance protocol (Fig 5A), flour replacement typically coincides with low abundance of larvae, and high abundance of pupae or young adults in the population. Hence, most of the larval inoculum might be effectively removed during the flour change, with very few remaining larvae that could contribute new fecal inoculum in the flour. Instead, as new adults eclose and mature, the flour accumulates increasing concentrations of quinones (18,19), which should suppress bacterial loads until the eventual increase in the number of larvae can counteract the effect of quinones. This plausible effect of flour replacement is supported by changes in the microbiome composition and total bacterial load in flour sampled across two generations in the same stock population (Fig 5B, Fig S17A-B). The disruptive effect of flour replacement is also seen in populations maintained under continuous generation cycles, where larval abundance does not fluctuate as much as under discrete generation cycles (Fig 5C, Fig S17C). In both sets of populations, bacterial loads dropped precipitously after a flour change, and gradually increased over time — coincident with increasing number of larvae — along with *Izhakiella* abundance.

**Fig 5:**
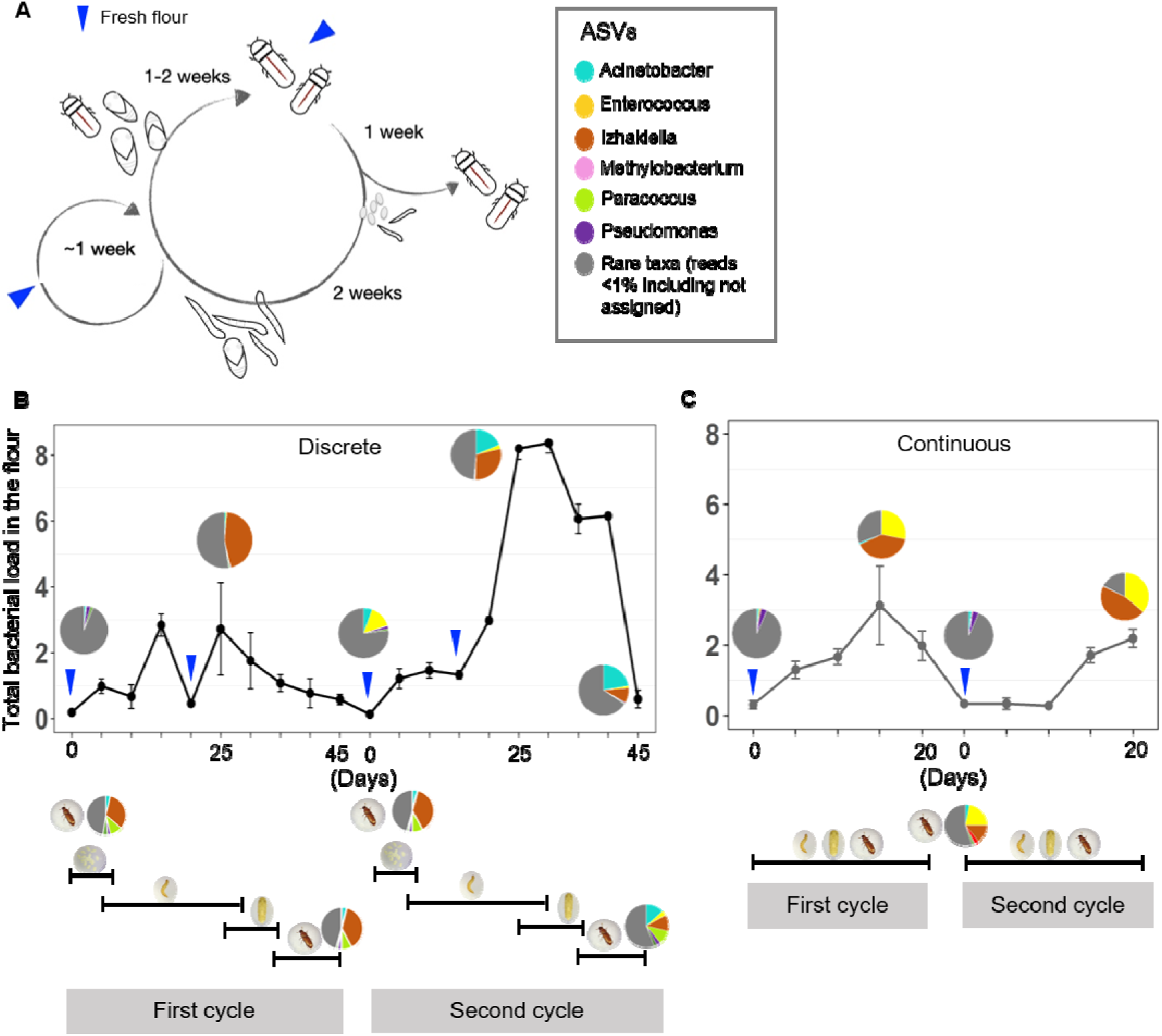
Effect of population maintenance protocol on the beetle microbiome. Blue arrows mark timepoints when the flour is replaced with fresh flour. (A) Schematic showing the stock maintenance protocol. (B) Results of Experiment 12 showing the time course of mean (±se) total bacterial load over time, in flour sampled every 5 days from stock populations maintained under discrete generation cycles (n = 2 technical replicates per timepoint). Pie charts show the average relative abundance of bacterial taxa in flour samples (within the plot area) and in sampled beetles (under the x-axis); the color key for the pie charts is placed next to panel A. Schematics along the x-axis indicate typical stage structure of the population at each timepoint. (C) Results of Experiment 12, showing data as described for panel B, for populations maintained under continuous generations. Fig S15 shows microbiome composition for each replicate in the flour as well as for beetles sampled for populations in both panels B and C.

Supporting our earlier results (Fig 1F), we again found that the total bacterial load strongly correlates with the relative abundance of *Izhakiella* in the flour from stock populations (Fig S16D, Spearman’s rank correlation rho = 0.5, P = 0.0001), reinforcing the conclusion that this genus is a strong colonizer of flour beetles. This is further evidenced by the mismatch in flour and beetle microbiomes (Fig 5B–C, beetle microbiomes shown under the x-axis; also see Fig S16 for individual level composition): at each sampling time, beetle microbiomes are often more enriched in *Izhakiella* than the flour microbiome.

### Microbiome has inconsistent effects on host fitness

The substantial microbiome variation observed above was surprising given that we had previously observed deleterious effects of disrupting the flour microbiome on several aspects of beetle fitness, especially for female fecundity (17) (Fig 6). We repeated these experiments over multiple years, finding that in most cases disrupting the microbiome of fresh flour using either UV treatment of flour or addition of antibiotics did not alter fecundity, in contrast to the previously reported results (Fig 6). In light of our current finding that fresh flour does not harbour significant bacterial loads, the lack of an effect of UV treatment is perhaps unsurprising. However, the lack of effect of antibiotics (which would kill bacteria growing inside the beetle gut) was more unexpected, and indicates the loss of beneficial bacteria after a point (Fig 6B), potentially due to microbiome drift. Finally, even when we added larval fecal inocula while rearing beetles from the egg stage, the emerging adults did not show a significantly higher fitness compared to adults reared without larval fecal input (Fig S6B). Together, these results show a lack of consistent fitness consequences of the beetle or flour microbiome on host fitness.

**Fig 6:**
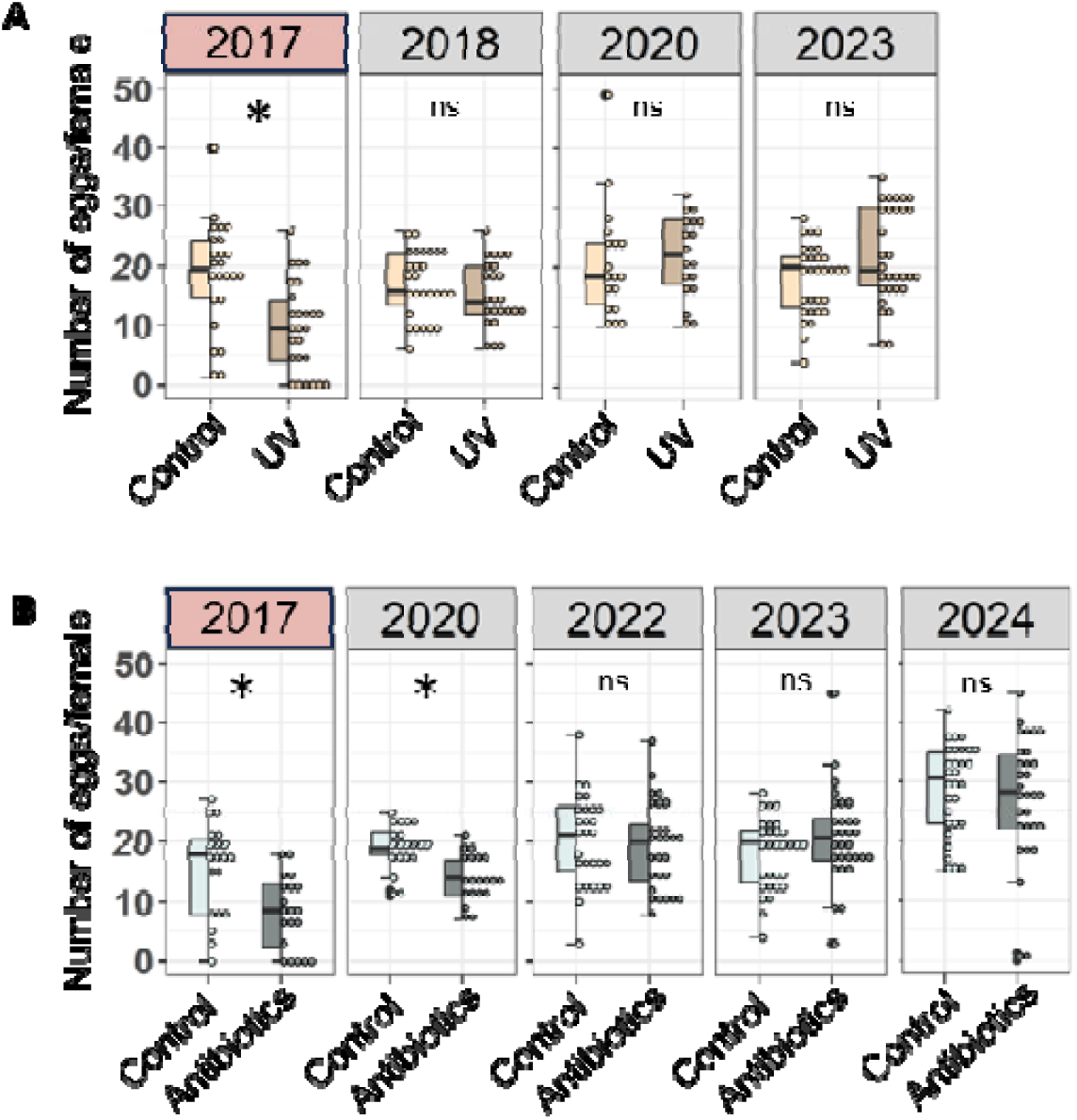
Inconsistent effects of depleting the flour microbiome on beetle fitness. (A) Results of Experiment 14, testing the effect of depleting the flour microbiome on female fecundity (n = 25-30 females/treatment/year). Fresh flour supplied to females for the assay was either untreated (control), or treated with (A) UV radiation or (B) a mix of three antibiotics. Asterisks indicate significant differences between control and test (t-test, p < 0.05). Pink facet headers represent data that were previously reported in (17), for comparison.

## DISCUSSION

Here, we identify and quantify the effect of multiple sources of stochasticity in community assembly and maintenance of the flour beetle microbiota. Our results show that several aspects of beetle life history and ecology explain the observed ecological drift in the microbiome (i.e., apparently idiosyncratic changes in the composition of the microbial community). (a) Bacteria are not maternally transmitted through the eggs or ovaries. (b) Larvae host large and consistent bacterial communities. Feces of larvae thus represent a key source of bacterial inoculum in the flour, likely because high larval feeding rates allow rapid proliferation and movement of bacteria between flour and gut, and between individuals. (c) In comparison, adult beetles harbor smaller and more variable microbiota. Hence, adult feces represent a smaller inoculum source, with the effect of lower feeding rates in adults compounded by the secretion of antimicrobial quinones. (d) Larval microbiomes stabilize rapidly compared to adult microbiomes, such that even a brief period of larval fecal enrichment can increase and homogenize flour and beetle microbial communities. (e) Stock maintenance protocols exacerbate ecological drift for two related reasons. First, the periodic replacement of used flour with fresh flour dramatically reduces flour microbial loads, mimicking frequent migration of adults in grain warehouses to colonize new resource patches. Second, at the time of such flour change, most larvae are either in the last instar (the non-feeding stage) or have already pupated, while newly eclosed adults begin secreting quinones. As a result, bacterial load in the flour does not increase rapidly. Drawing upon these results, we propose a model for bacterial transmission, persistence and host colonization in the flour beetle system (Fig 7). Together, the periodic flour replacement and absence of larval fecal inoculum, lack of vertical bacterial transmission, and quinone secretion by adults cause substantial stochasticity in community assembly, leading to the observed ecological drift in adult flour beetle microbiomes. Flour beetles may thus serve as a good example of neutral community assembly, with a short host lifespan and weak microbial dispersal into hosts: two factors that are predicted to drive large individual variation in microbiome composition (13). Our results contribute to the growing body of research on the role of stochastic colonization on insect-microbe associations, which is important to more broadly understand insect populations and adaptation to environmental change (7).

**Fig 7:**
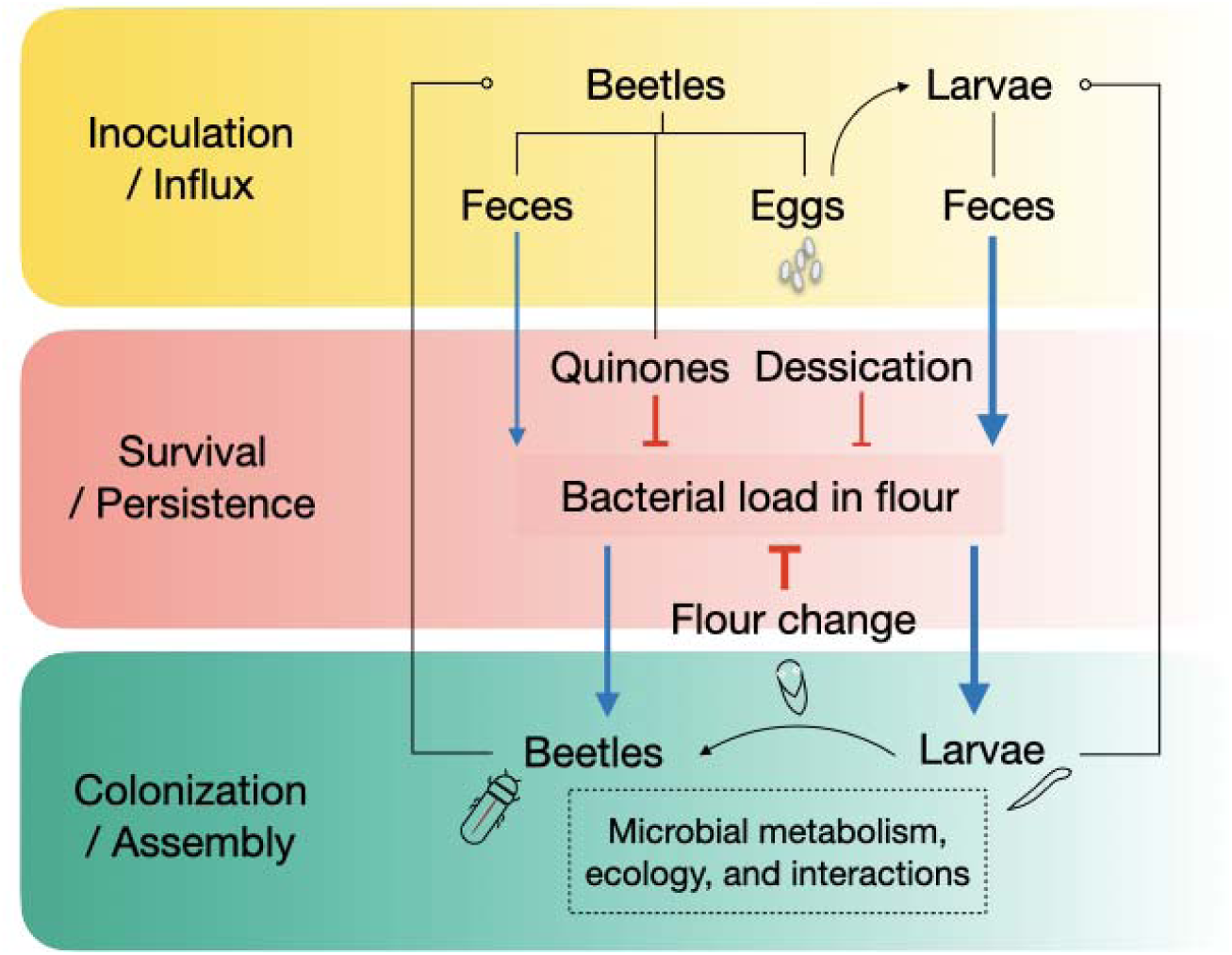
Proposed model for key processes shaping the red flour beetle microbiome. We consider three different aspects of host-bacterial interactions: bacterial inoculum and survival in the flour, and host colonization. Life cycle transitions are shown with curved arrows. Blue arrows indicate increased bacterial transmission; red indicates inhibitory effects; increasing arrow thickness indicates stronger effects. Hypothesized factors with unknown effects (not tested in this study) are marked by a box with dashed borders.

We suggest that our inability to replicate our prior results showing significant fitness benefits of beetle-associated microbiomes (17) may arise due to microbiome drift in our stock populations over time. One possibility is that the wild-collected founder beetles harboured beneficial bacteria that were gradually lost after we established laboratory stocks in 2013, exacerbated by ecological drift due to periodic population maintenance protocols. However, in the wild, adult beetles routinely disperse and colonize fresh flour patches (36), a situation similar to flour replacement in the laboratory that should also lead to significant drift. Further, this scenario cannot explain why we observed a significant decline in fitness after treating fresh flour (with very low microbial loads) with UV radiation, including not only reduced fecundity but also reduced survival and lifespan (17). An alternative explanation is that some batches of fresh flour used before 2017 served as a source of beneficial bacteria that were missing in subsequent batches, explaining both the significant deleterious effects of UV and antibiotic treatment until 2017-2018, and the subsequent lack of such effects. Deeper analysis of beetle microbiomes over the years may provide some answers. However, regardless of the exact cause for initially highly repeatable and later irreproducible fitness effects, it is clear that the effects of microbiota are not consistent over time, coincident with large ecological drift in the microbiome. It is plausible that the chance loss of putative beneficial bacteria may have allowed and further exacerbated this drift over time; that the benefit provided by the bacteria was not sufficiently large to facilitate rapid evolution of transmission mechanisms, in turn preventing the overpowering effect of drift; that we happened to catch flour beetle populations in a transient state where a potential mutualism failed to establish; that selection for traits relevant for adaptation to laboratory conditions resulted in the dissociation of the host-microbiome relationship; or some combination of or all of the above. These are exciting open questions.

More generally, our work allows us to speculate about the mechanisms that may prevent or facilitate the establishment of obligate host-microbial interactions, which involve both host and bacterial control (37). A recent mathematical model showed that in the absence of strict vertical transmission, horizontal or environmental transmission of microbiota is critical for the establishment of a host-microbial relationship (38). In *T. castaneum,* we did not observe evidence of vertical transmission, and we find that horizontal transmission is context-dependent. The lack of vertical transmission may be interpreted in different ways. It may arise due to constraints placed by ecological drift in adult microbiomes, e.g., because young ovipositing females do not have a stable microbiome to transmit. In turn, the absence of such transmission may also contribute towards preventing the establishment of a stable microbiome, by enhancing microbiome stochasticity at the beginning of an individual’s development. Thus, the lack of vertical transmission may explain why a strict host-microbe relationship has not evolved in flour beetles, despite evidence for host fitness benefits (17). On the other hand, weak (rather than strong) vertical transmission could also be selectively favoured, e.g., because it can drive among-host variation that is adaptive in fluctuating environments (39). It is plausible that the frequent colonization of new habitat patches by *T. castaneum* — a feature of its life history as a human-associated pest — may generate such selection for weak vertical transmission of microbiota. Yet another possibility is that strong selection on quinones as signals of population density — important for negative density dependent population regulation (34) — may also indirectly impede the establishment of a stable microbiome. Finally, it is possible that all of these mechanisms contribute to the lack of stable host-microbiome associations in flour beetles, and that similar processes may be at play in several other species that lack such relationships (6). Although distinguishing cause and effect in our case is difficult, it may be possible via evolution experiments where each contributing factor can be independently manipulated and its effect on the emergence of vertical transmission quantified. Prior laboratory experiments with stink bugs suggest that such evolution is plausible within relatively short timescales (40). More generally, experimental evolution approaches may be fruitful to understand the relative contribution of several factors driving stochastic vs. deterministic dynamics, and the emergence of strong host-microbial associations or lack thereof.

Our results also complement prior work on microbiome variation in insects that undergo large ecological shifts across developmental stages (41). Dramatic internal reorganization during metamorphosis in holometabolous insects (including beetles) is associated with high beta diversity and microbiome turnover, compared to hemimetabolous insects (42). Microbiome shifts across life stages are also common in insects (43), including dragonflies ((44), butterflies (4,45), and mosquitoes (46). For instance, gut microbiome assembly in monarch butterflies mirrors the patterns we observed in flour beetles, whereby the microbiome load is high in larvae and decreases in pupae and adults (45). However, in most cases, the microbiome shift across insect life stages is typically associated with a niche shift that can directly influence the microbiome by altering the influx of bacteria and/or available nutrients. The case of *T. castaneum* is remarkable because the microbiome shift is not associated with a diet or habitat change; in fact all life stages share the same habitat that serves as the route for rapid fecal-oral transmission. We speculate that in flour beetles, the life stage-specific variation may derive from differential feeding rates (allowing, e.g., faster bacterial proliferation in larval guts as reported in (47,48), distinct gut morphology (e.g., the larval gut may be more suitable for bacterial growth(43), or different immune responses (e.g., adults may have more effective immune function, suppressing bacterial growth in the gut (49). Note that quinone secretion by adults probably does not contribute to stage-specific microbiome variation, because quinones suppress overall and taxon-specific microbial load in the flour, which should therefore influence available inocula for all life stages. We hope that future work can distinguish between these causes of life stage-specific variation in flour beetle microbiomes.

Here, we analysed broad patterns of variation in microbiota composition, demonstrating significant ecological drift. However, this does not imply that community assembly is entirely stochastic: indeed, we observed that a relatively small number of genera (such as *Izhakiella* and *Enterococcus*) repeatedly dominated, indicating a role for deterministic factors. Thus, several questions remain about factors that guide community assembly and maintenance — which specific taxa are more vs. less likely to colonize, and under what conditions? In the fruit fly, successful colonization by a focal species depended on stochasticity as well as the identity and presence of other species, likely due to deterministic outcomes of inter-species interactions (50). Similarly, processes and phenomena such as priority effects, differential survival, differential competitive ability, and niche use may determine precisely which taxa will dominate the beetle microbiome at a given time (Fig 7). In our case, it remains unclear whether, why, and when *Izhakiella* is a strong colonizer; this requires further work, which is currently challenging due to our inability to culture the beetle-associated strain of *Izhakiella*. We also acknowledge that our current analysis is coarse-grained, and further interesting patterns may emerge at the level of variation and dynamics across bacterial strains, or at higher level functional classes. We hope to address these issues in future work. Finally, in closing we note that even non-essential bacterial symbionts may significantly influence host ecology and life history, e.g., by providing transient benefits such as protection against parasites, predators, and pathogens (Łukasik & Kolasa, 2024)(Łukasik & Kolasa, 2024). It remains to be seen whether any of the observed colonizers of the flour beetle gut provide the host with fitness benefits, or whether the community truly represents neutral assembly.

## ACKNOWLEDGEMENTS

We thank Basabi Bagchi and other members of the Agashe lab for discussion and critical comments on the manuscript; Prapti Satpathy, Manjunath Reddy, Soumya Panyam, and Gaurav Agavekar for laboratory assistance, and Sampath Kumar and the insectary facility at the Tata Institute for Genetics and Society for training on and access to their microinjection platform. We acknowledge funding and support from the National Centre for Biological Sciences and the Department of Atomic Energy, Government of India (Project Identification No. RTI 4006); the DBT/Wellcome Trust India Alliance (award number IA/I/17/1/503091 to DA); and an SERB National Postdoctoral Fellowship (PDF/2018/001441 to RD).

## AUTHOR CONTRIBUTIONS

DA, PS, RD, SG, SS, SB designed the project; all authors conducted experiments; PS analysed data; DA and PS wrote the manuscript with input from all authors; DA acquired funding.

## DATA AVAILABILITY

All the raw sequencing data are available in the Sequence Read Archive (SRA) of the National Centre for Biotechnology Information (NCBI) under Bioproject (PRJNA1330605). 16S rRNA Sanger sequenced data are available in the GenBank of the National Centre for Biotechnology Information (NCBI) with accession numbers (PX376855-PX376865, PX377900, PX377901). All processed data used for analysis are available on Figshare (10.6084/m9.figshare.30135973).

