## Supplemental info for "Ecological drift in the flour beetle microbiome and associated instability in host benefits"

**SUPPLEMENTARY TABLES**

**Table S1: List of bacterial strains isolated from beetles and screened for sensitivity to quinones.** Media used: M–Enterococcus Agar (HiMedia), Nutrient agar (HiMedia) with 0.5% wheat bran added in some cases; R2A agar (Reasoner’s 2A Agar, HiMedia) supplemented with 15% or 30% sucrose; TSA (Tryptic Soy Agar, HiMedia), BHI (Brain Heart Infusion Agar, HiMedia), Urea Base Agar (HiMedia), and Luria Agar (Difco). All conditioned flours were sampled directly from the stock population boxes. The number inside the parentheses indicates the number of samples used for the isolation process.

| Genbank accession no | 16S rRNA identification | Media used | Source |
| --- | --- | --- | --- |
| PQ432978 | *Enterococcus faecium* | M – Enterococcus Agar | Crushed adult male beetle (3) |
| PX376859 | *Serratia nematodiphila* | Nutrient Agar | Adult dissected gut (10) |
| PQ436511 | *Pseudomonas mosselii* | Nutrient Agar | Adult female dissected gut (10) |
| PQ433218 | *Acinetobacter radioresistens* | Nutrient Agar + wheat bran | Conditioned wheat flour |
| PX377900 | *Acinetobacter baumanni* | R2A + 30%sucrose | Crushed adult male beetles (3) |
| PX377901 | *Acinetobacter lwoffii* | R2A + 30%sucrose | Crushed adult female beetles (3) |
| PX376860 | *Aureimonas* | Nutrient Agar + wheat bran | Adult dissected gut (10) |
| PQ432977 | *Enterococcus faecalis* | M – Enterococcus Agar | Conditioned wheat flour |
| PRJNA33175 | *Izhakiella capsodis* | R2A + 15%sucrose+3%NaCl | Belgian Coordinated Collections of Microorganisms (BCCM), strain described in Aizenberg-Gershtein et al 2016 |
| PX376861 | *Aeromonas dhakensis* | TSA | Adult dissected gut (10) |
| PX376862 | *Aeromonas caviae* | TSA | Adult dissected gut (10) |
| PX376863 | *Enterobacter cloacae* | TSA | Adult dissected gut (10) |
| PX376864 | *Pseudescherichia vulneris* | BHI | Live adult beetles (8) |
| PQ432976 | *Enterobacter hormaechei* | BHI | Live adult beetles (8) |
| PQ432980 | *Staphylococcus gallinarum* | BHI | Conditioned wheat flour |
| PX376865 | *Mixta calida* | BHI | Conditioned wheat flour |
| PX376858 | *B thuringiensis* | Urea base agar | Crushed 14-20^th^ days old larvae (50) |
| PX376857 | *Pseudomonas parafulva* | Urea base agar | Crushed 14-20 days old larvae (50) |
| PX376856 | *Stutzerimonas stutzeri* | Urea base agar | Crushed 14-20^th^ days old larvae (50) |
| PX376855 | *Cronobacter sakazaki* | Luria Agar | Crushed 14-20^th^ days old larvae (50) |
| PQ432979 | *Planococcus sp.* | R2A + 15%sucrose+3%NaCl | Conditioned wheat flour |

**Table S2: Effect of treatment (with vs. without quinones) and time on flour microbial communities, PERMANOVA.** Significant P-values are shown in bold.

|  | Treatment | | Time | | Treatment x time | |
| --- | --- | --- | --- | --- | --- | --- |
|  | R^2^ | P | R^2^ | P | R^2^ | P |
| Fresh flour | 0.03 | 0.77 | 0.12 | **0.018** | 0.03 | 0.95 |
| Fresh flour +*Acinetobacter* | 0.048 | 0.77 | 0.12 | **0.013** | 0.048 | 0.76 |
| Fresh flour + *Enterococcus* | 0.33 | 0.88 | 0.17 | **0.0009** | 0.03 | 0.89 |
| Fresh flour + adult feces | 0.023 | 0.98 | 0.12 | **0.037** | 0.023 | 0.98 |

**SUPPLEMENTARY FIGURES**

**Fig S1: Schematics showing the design of Experiments 1a, 1c and 2.** (A) Experiment 1a compares adult female microbiomes across time, from the same stock population. (B) Experiment 1c compares adult female microbiomes from independent stock population boxes (founded by the same parental stock), sampled at the same time. (C) Experiment 2 compares microbiomes of adult offspring across 3 replicate boxes set up using the same stock parents.

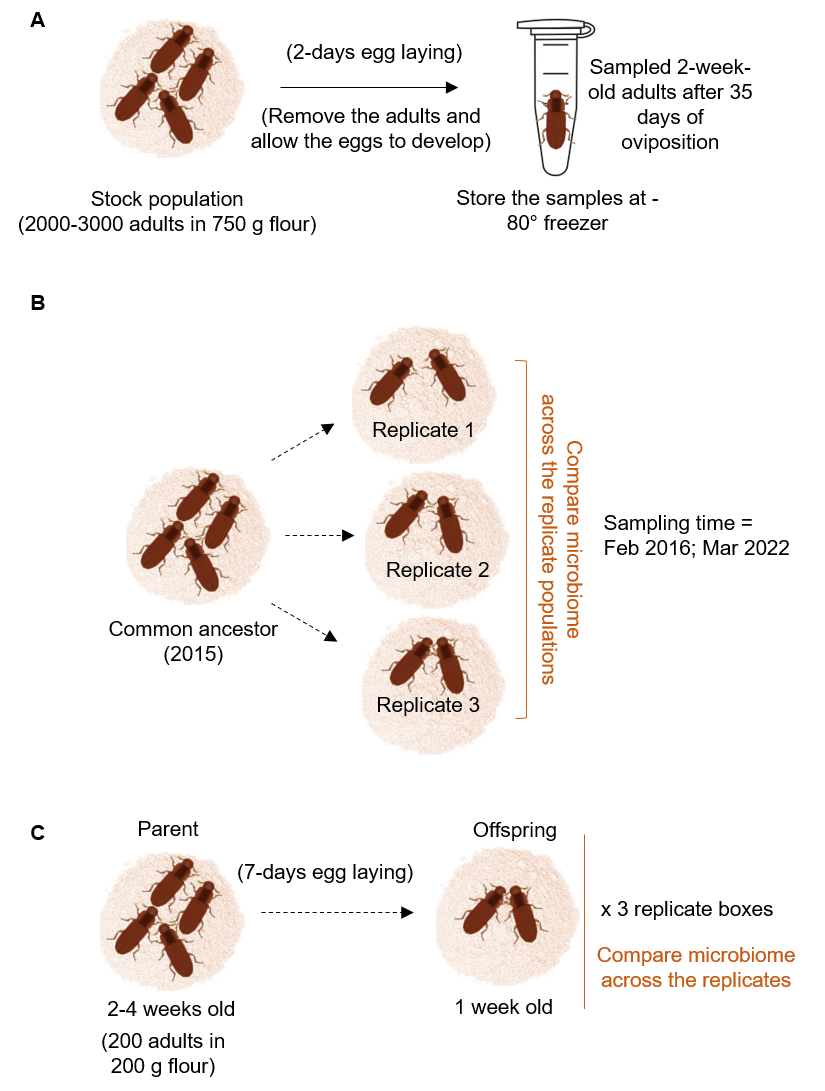

**Fig S2: Schematics showing the design of experiments to test the effect of external environmental factors and adult density.** (A) Experiment 3 compares the microbiomes of beetles reared under different treatment conditions. For air sterility and flour treatment, we used we used 2 days of egg laying; for experimenter effect, we used 7 days of egg. (B) Experiment 5 compares the microbiomes of the 2-week-old adults set up across two different adult densities (LD = low density, HD = high density).

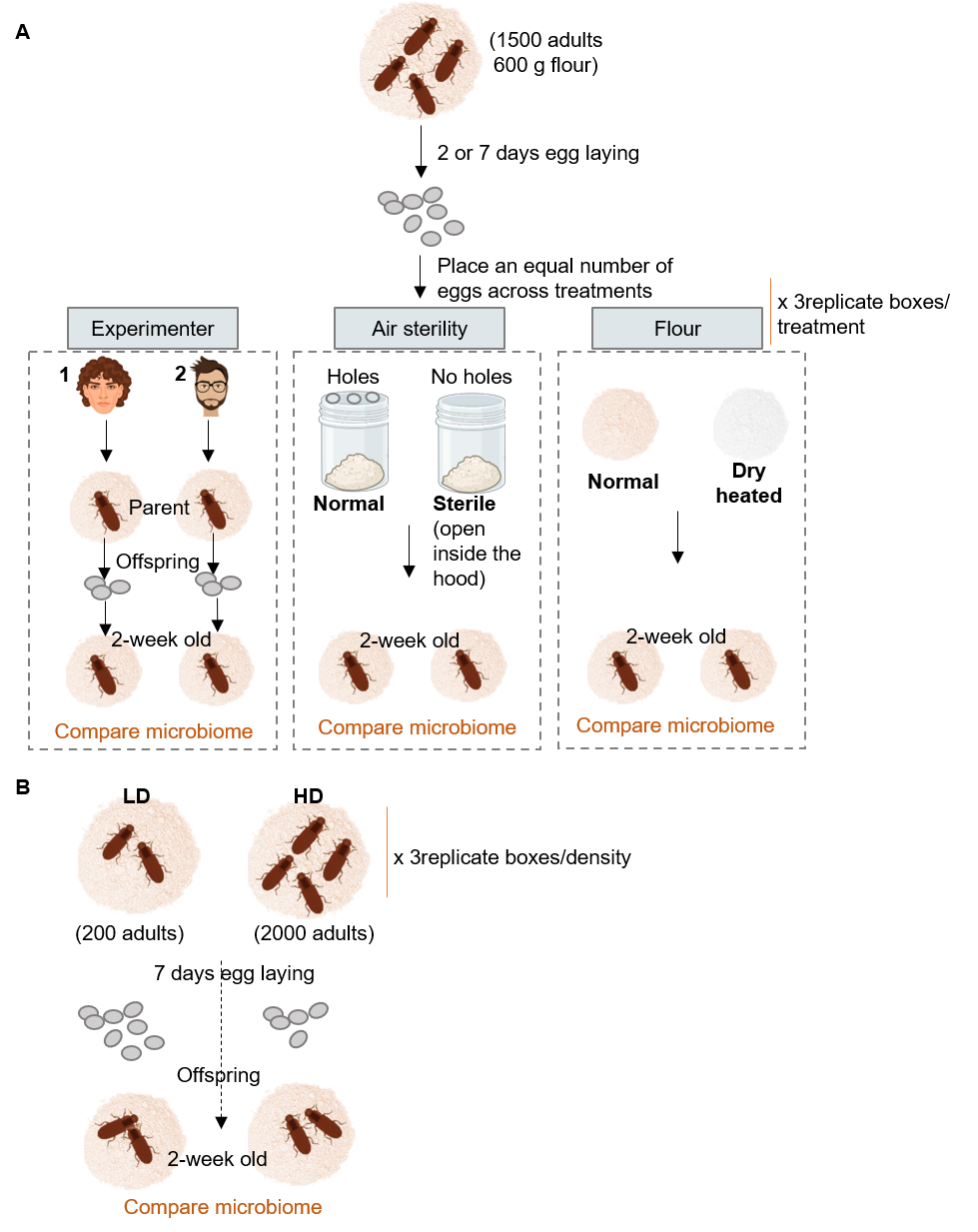

**Fig S3: Schematics showing the design of experiments to test for maternal microbiome transmission to eggs.** (A) Experiment 4a compares the microbiomes of mothers and pooled eggs laid in flour. (B) Experiment 4b compares microbiomes of eggs and ovaries dissected from replicate females. (C) Experiment 4c compares the microbiomes of eggs dissected from females of different ages.

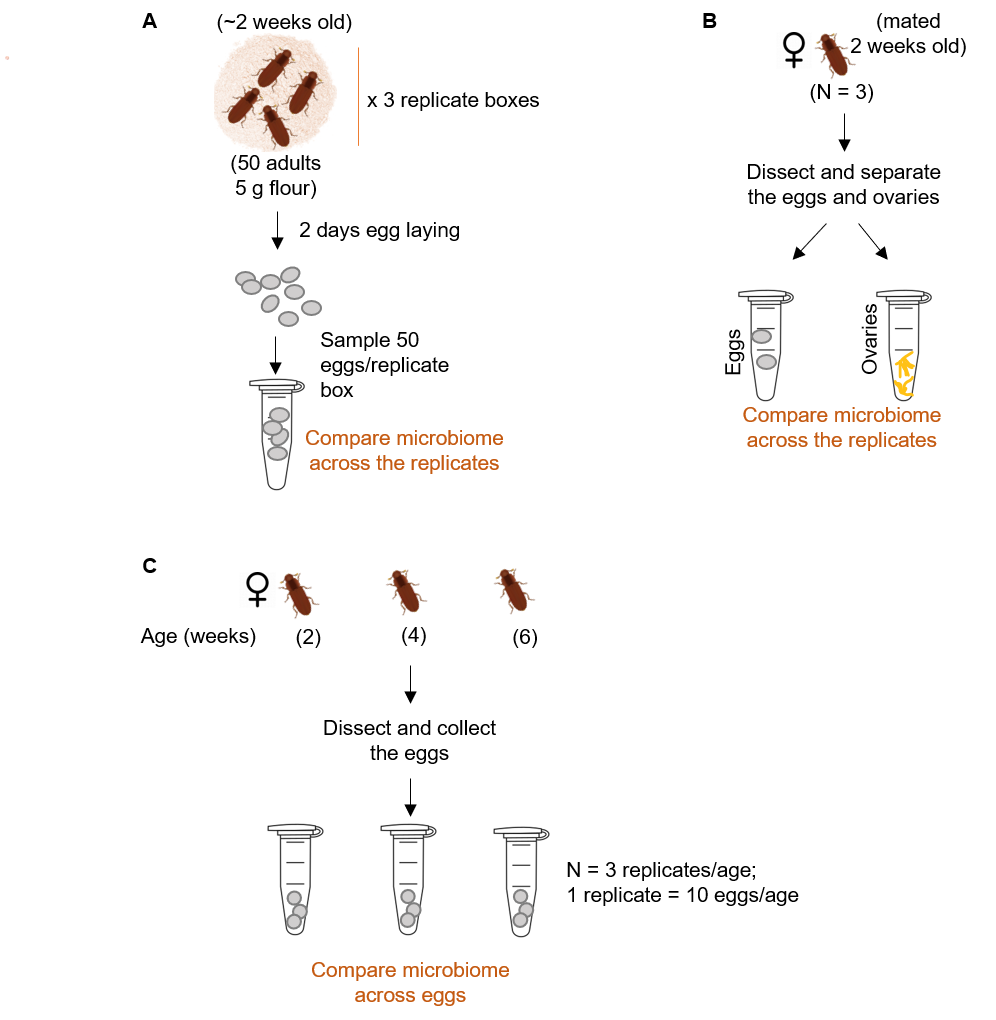

**Fig S4: Schematics showing the design of experiments to test the effect of feces on microbiomes.** (A) Experiment 6 compares the microbiota of feces collected from larvae and adults across development. (B) Experiment 7 compares the microbiomes of adults fed with either fresh flour, or flour enriched with larval or adult feces, or flour with both larval feces and adult stink gland extract. (C) Experiment 8 compares microbiomes of adults fed with flour containing different proportions of larval feces.

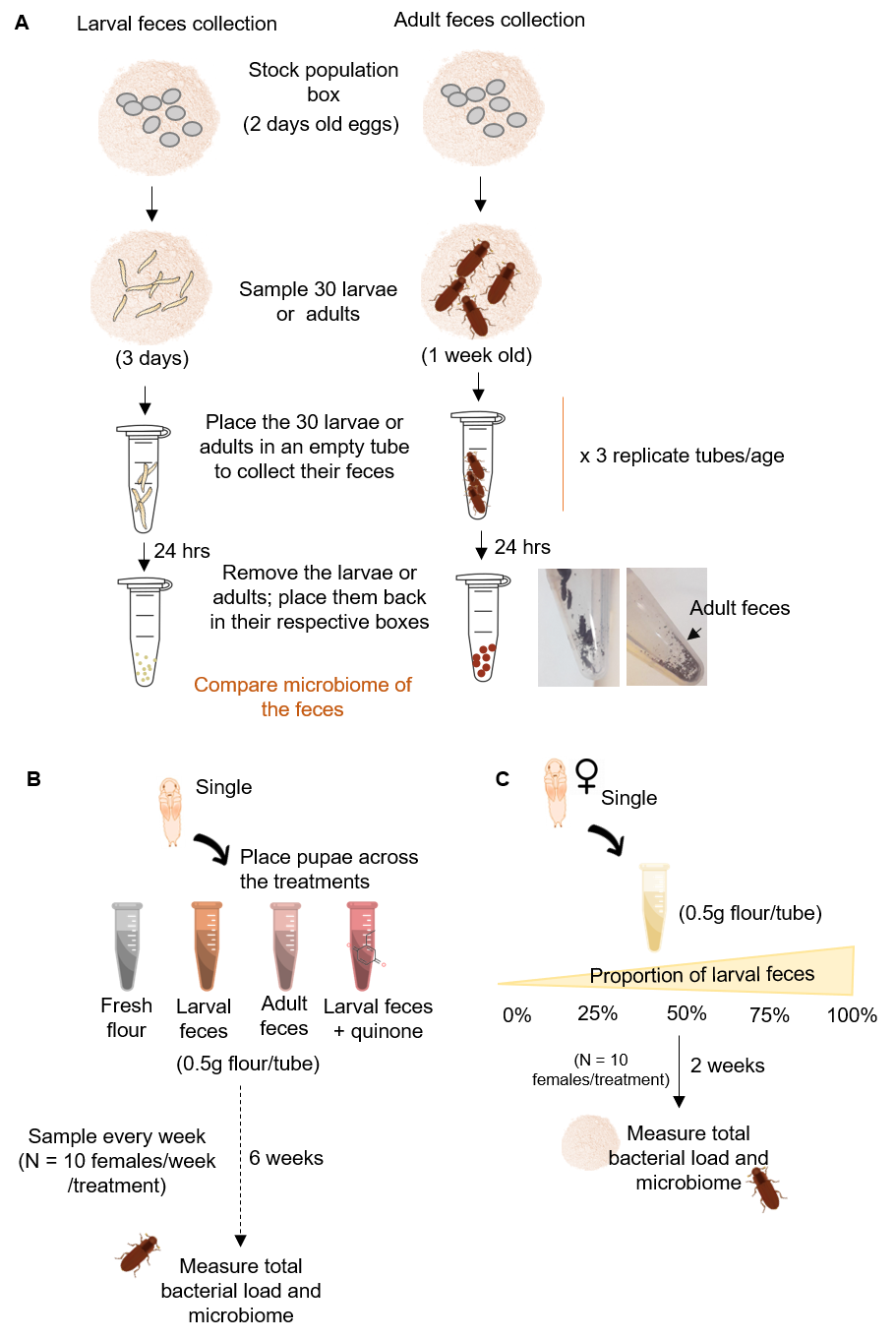

**Fig S5: Schematics showing the design of experiments to test the effect of quinones on the beetle microbiome.** (A) Experiment 10 compares the microbiomes of the flour collected every 4^th^ day of the treatments. (B) Experiment 11 compares the microbiome of adults with RNAi-mediated knockdown of GFP (negative control) or GT39 (quinone synthesis). Two control treatments included no injection (untreated), or injection with buffer alone. (C) Dissected beetles confirming the absence of stink glands in the dsGT39-treated beetles.

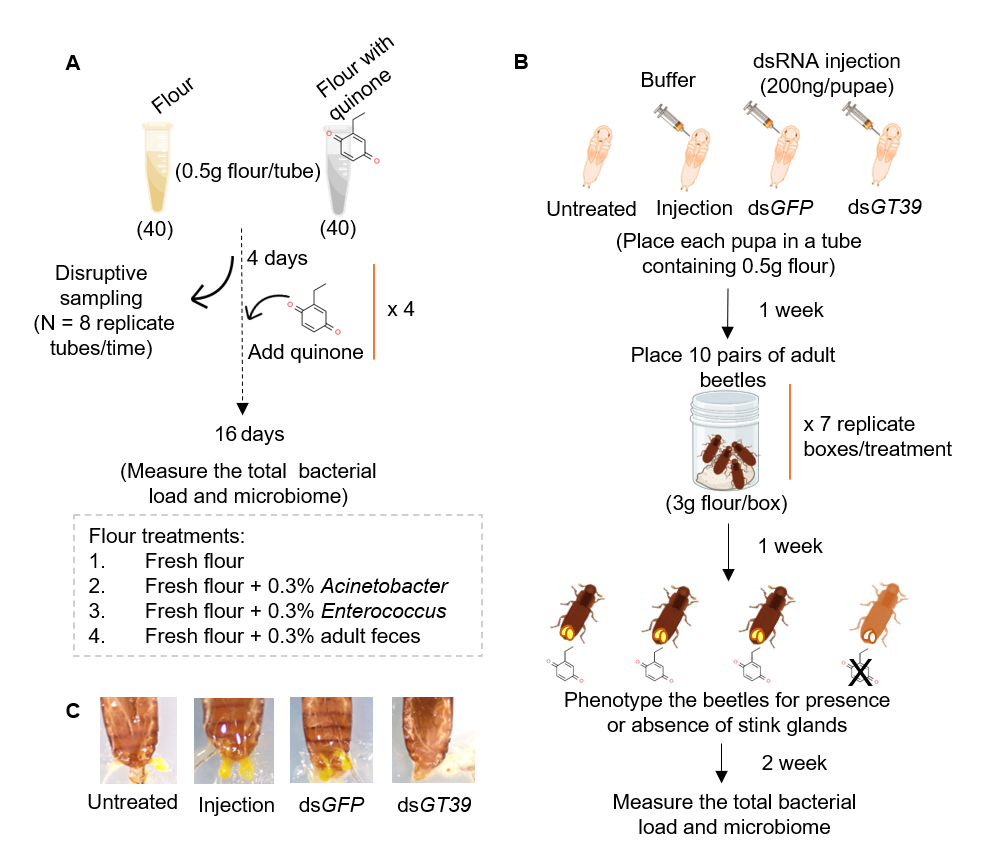

**Fig S6: Measuring the effect of microbiome on beetle fitness.** (A) Design of Experiment 13b, showing the effects of adding larval feces (inoculum) on beetle fitness. W = fresh wheat flour, ENL = mix of 50% fresh wheat flour and 50% flour enriched with larval feces (see Methods for Experiment 7). (B) Results from Experiment 13b. Each point represents total offspring produced by a female. For all pairwise comparisons across treatments (t test), P >0.05.

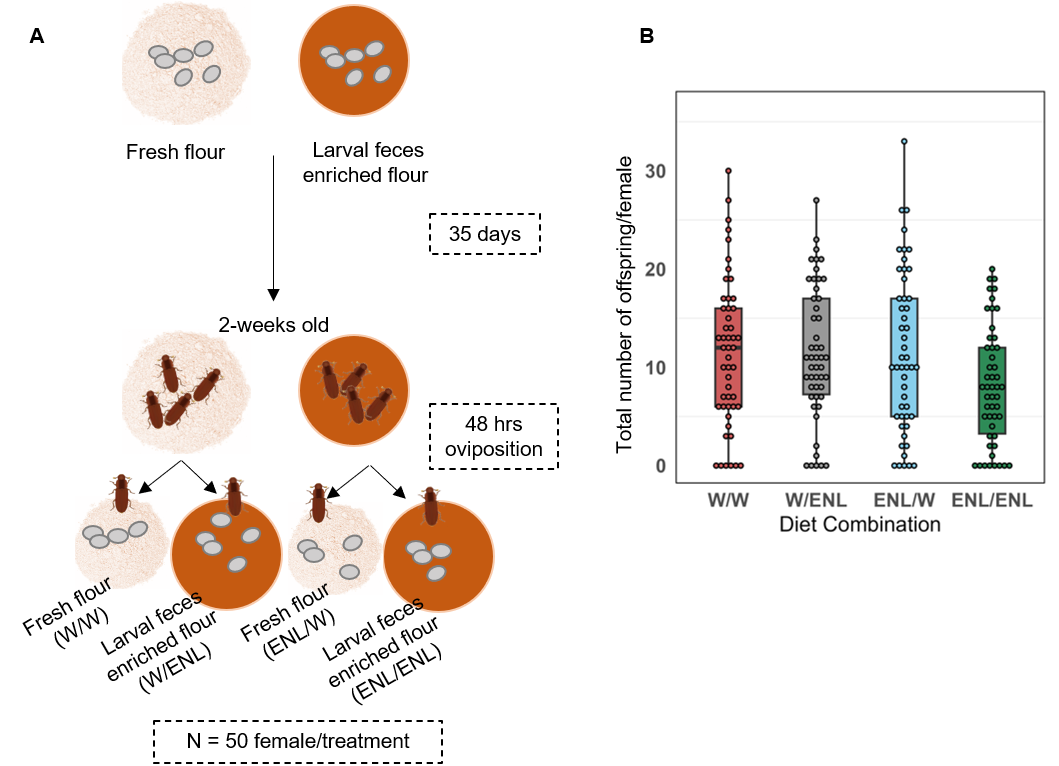

**Fig S7: Rarefaction plots show the number of ASVs (richness) as a function of the number of reads sampled from beetle and wheat flour samples.** Each line represents a sample, and error bars represent the standard error computed from 1000 random subsamples from the total reads. In both cases, we chose samples with the maximum reads (eg. 50000 reads in beetle samples; and 10000 in flour samples). We used 6 beetle samples from dataset of Fig 1A, and flour samples from multiple experiments. Dotted grey lines represent the threshold we set for the minimum number of reads per sample required for further analysis.

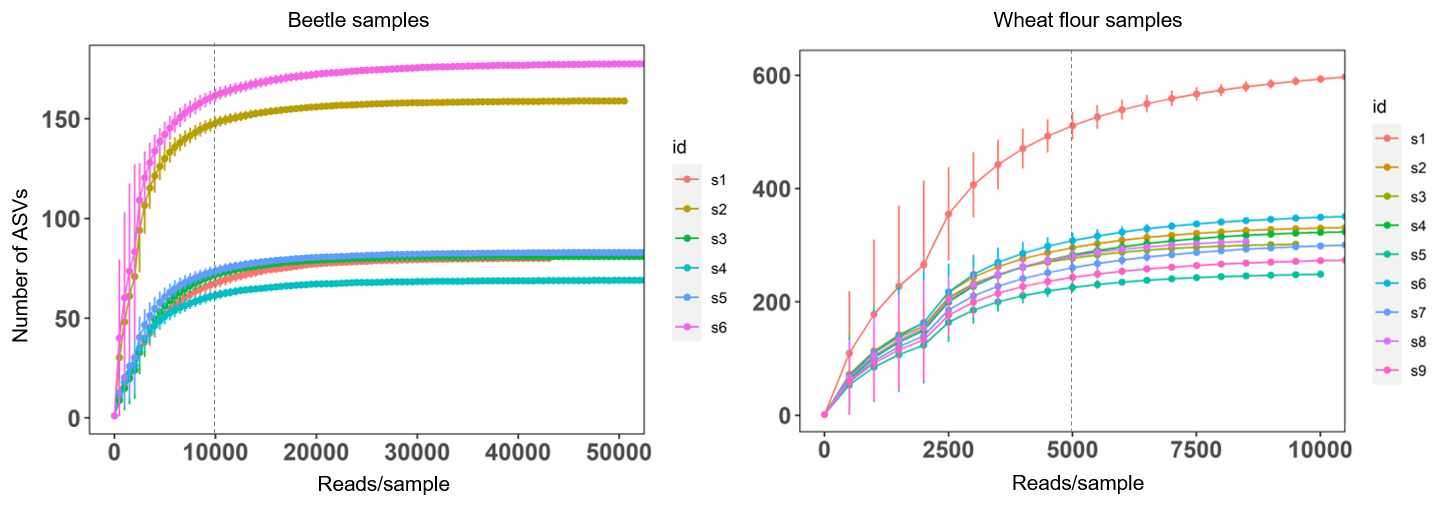

**Fig S8: Bacterial community richness and diversity in female beetles sampled across multiple months and years.** Violin plots show (A) the number of ASVs (richness) and (B) Shannon diversity index, for data shown in Fig 1A. Sample sizes are given in Fig 1A.

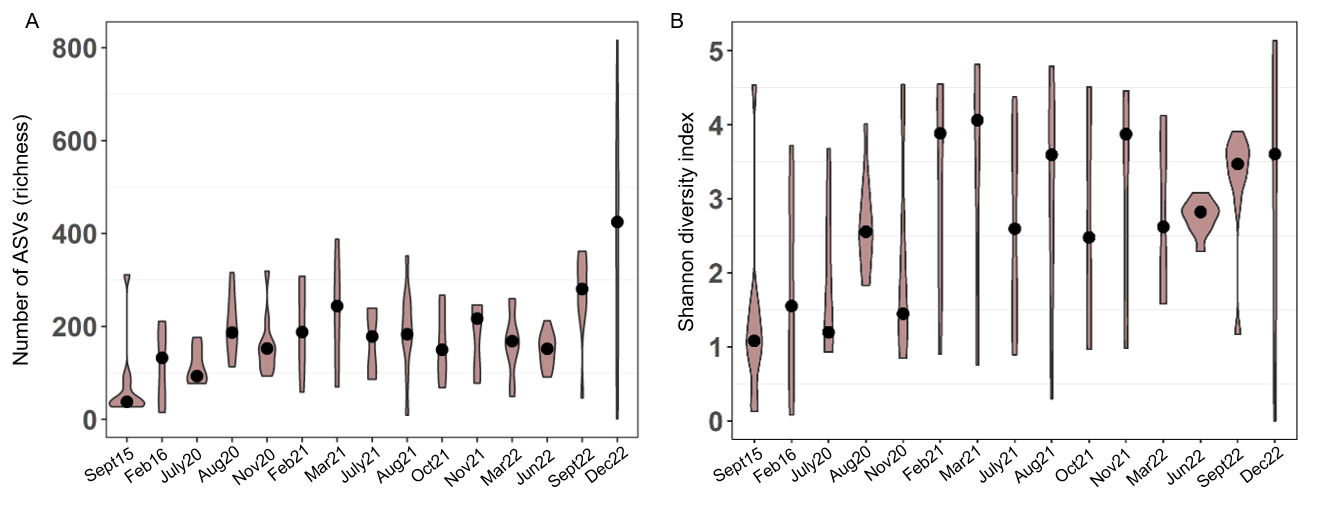

**Fig S9: Microbiome variation in male beetles.** Stacked bar plots show the relative abundance of bacterial ASVs (identified to genus level) for each male beetle sampled from stock populations (Jul 2020, Feb 2021) and three independent stock lineages in Mar 2022. Sample size (number of male beetles) is indicated in parentheses.

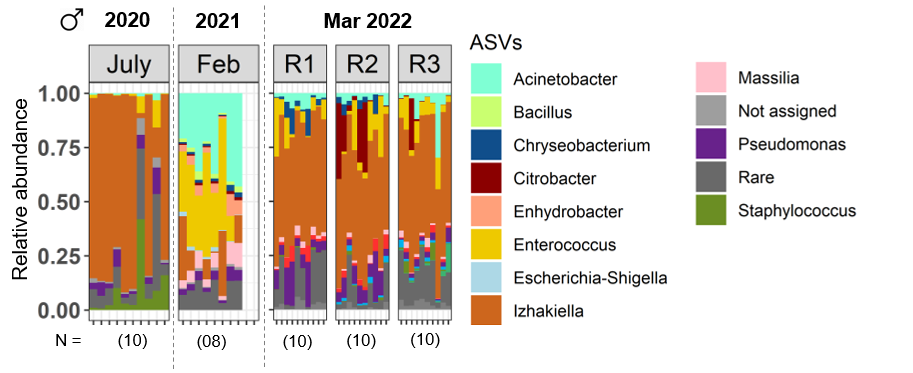

**Fig S10: Core microbiome of the red flour beetle**. The heatmap shows the prevalence of each bacterial taxon across 146 beetles (from data shown in Fig 1A) at different relative abundance thresholds. Prevalence of the taxon is calculated as the proportion of samples containing that taxon at or beyond the indicated relative abundance threshold. E.g., *Izhakiella* is prevalent in most (80%) samples across the range of relative abundances of 0.01-0.1, whereas *Acinetobacter* is prevalent (50%) only at low abundance (0.01).

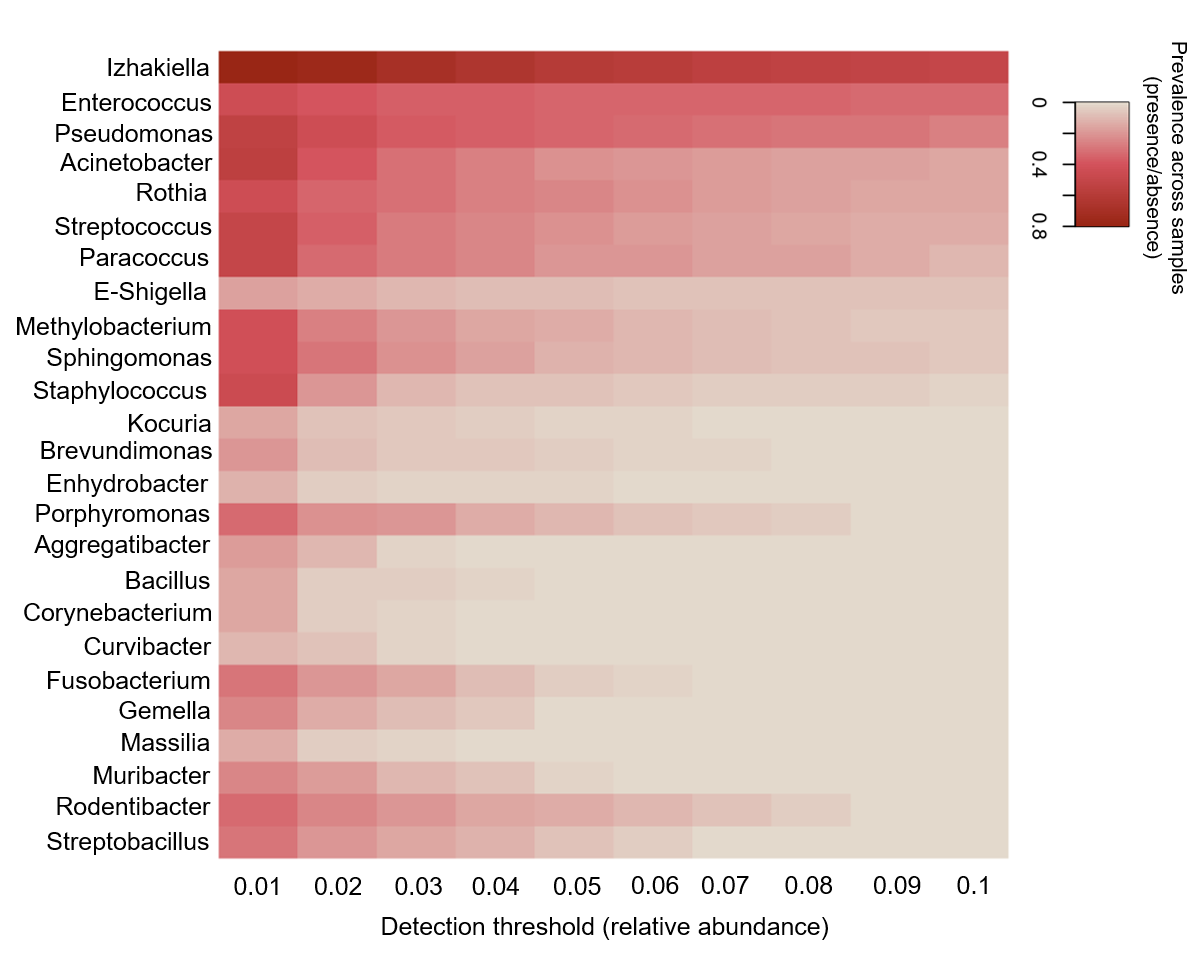

**Fig S11**: **The total bacterial load is not correlated with the relative abundance of *Enterococcus* or other, rare taxa.** Scatterplots show the relationship between the normalized total bacterial load and the relative abundance of *Enterococcus* or ‘other’ taxa (combining reads of rare taxa and those with unassigned taxonomy), for beetle samples showed in Fig 1E. Results of non-parametric correlation tests are shown. Lines indicate best fit regression with confidence intervals.

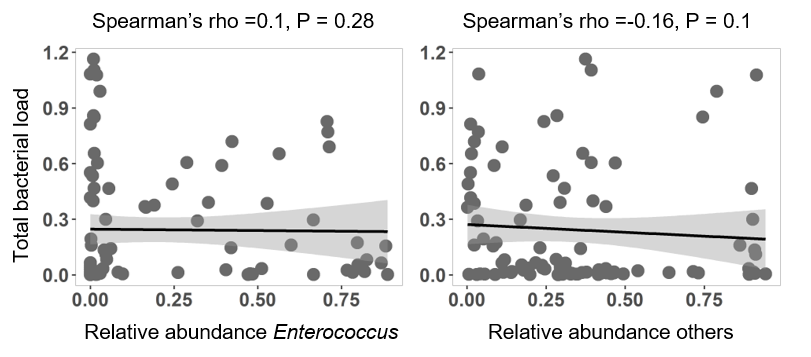

**Fig S12: External environmental factors explain very little variation in the beetle microbiome.** Results of Experiment 3, testing the effects of external factors. (A) Bacterial community composition in female beetles as a function of experimenter identity, air sterility, and flour treatment; each factor was tested in separate experiments. (B) Bacterial community composition in adult males. (C) Taxonomic beta diversity (β_rc_) of bacterial communities for samples shown in panel A. (D) Mean total bacterial load (normalized to total host DNA) for a subset of samples shown in panel A (n = 3 females/treatment). Error bars represent standard error. For reference, the dotted grey line indicates the mean total bacterial load per beetle observed for the samples shown in Fig 1E.

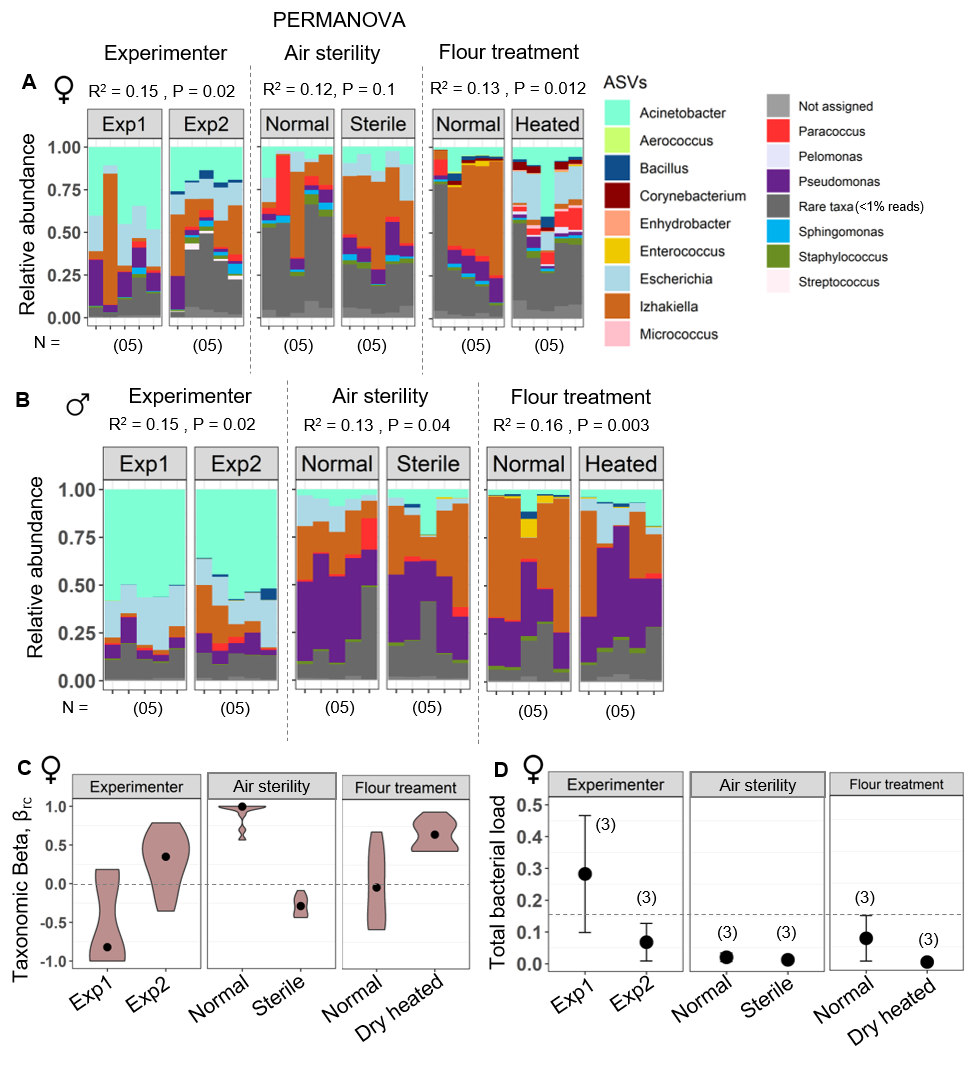

**Fig S13:** **No evidence for consistent maternal microbiome transmission to eggs.** (A) Results from Experiment 4a, determining bacterial community composition in 2-week-old mothers and their eggs. (B) Image showing an example of dissected ovaries and eggs from a 2-week-old female. (C) Results of Experiment 4b, determining bacterial community composition in ovaries dissected from 2-week-old females (O) and their eggs (E). (D) Results of Experiment 4c determining the bacterial community composition in eggs dissected from ovaries of females of different ages. Sample sizes are indicated in parentheses below bars.

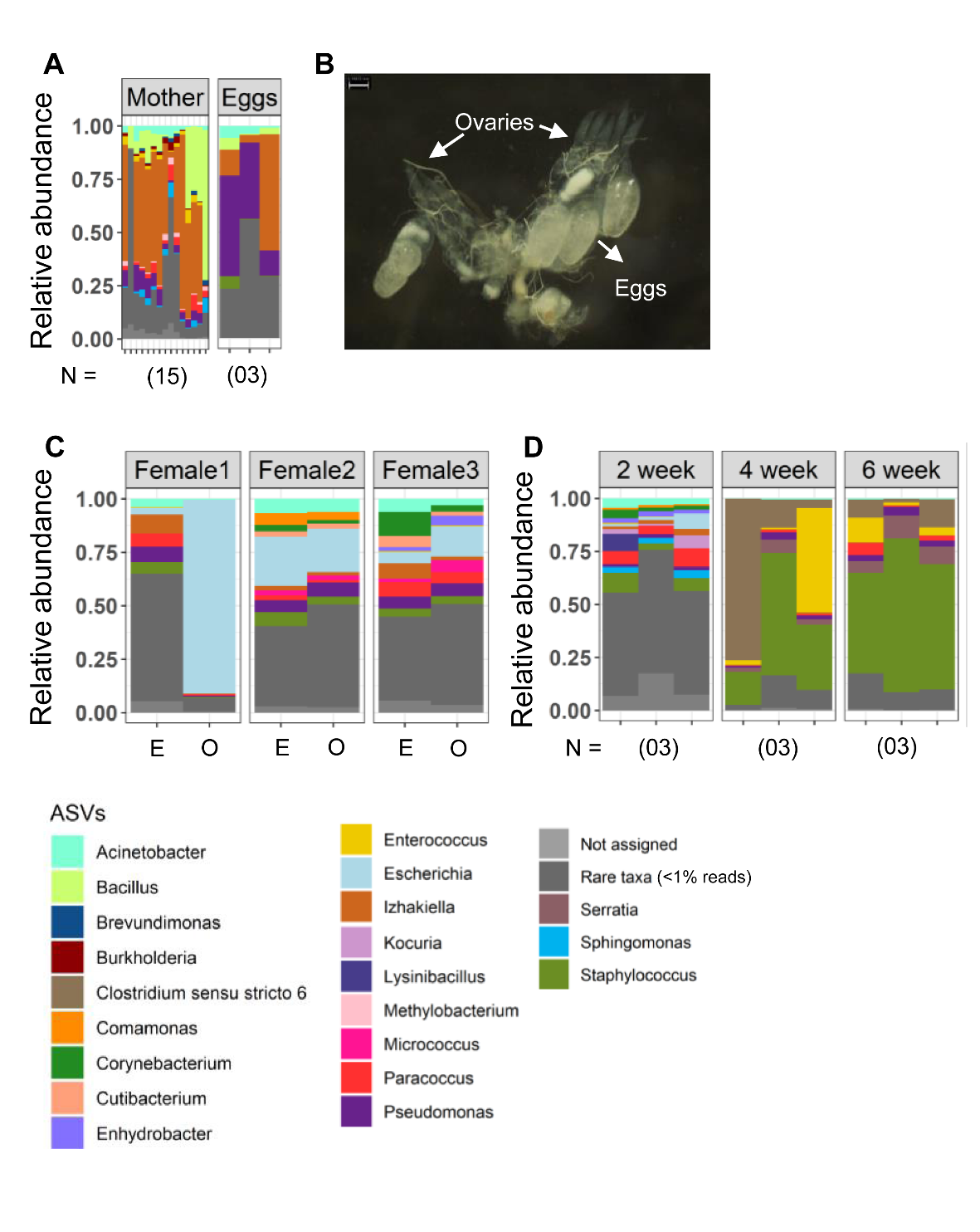

**Fig S14: Microbiome variation across males reared under varying adult parent densities.** Stacked bar plots show bacterial community composition across 2-week-old males that developed under varying adult densities (LD = low density of 200 adults; HD = high density of 2000 adults). R1-R3 indicate replicate population boxes created from the same parental stock. Sample size (number of male beetles) is indicated in parentheses.

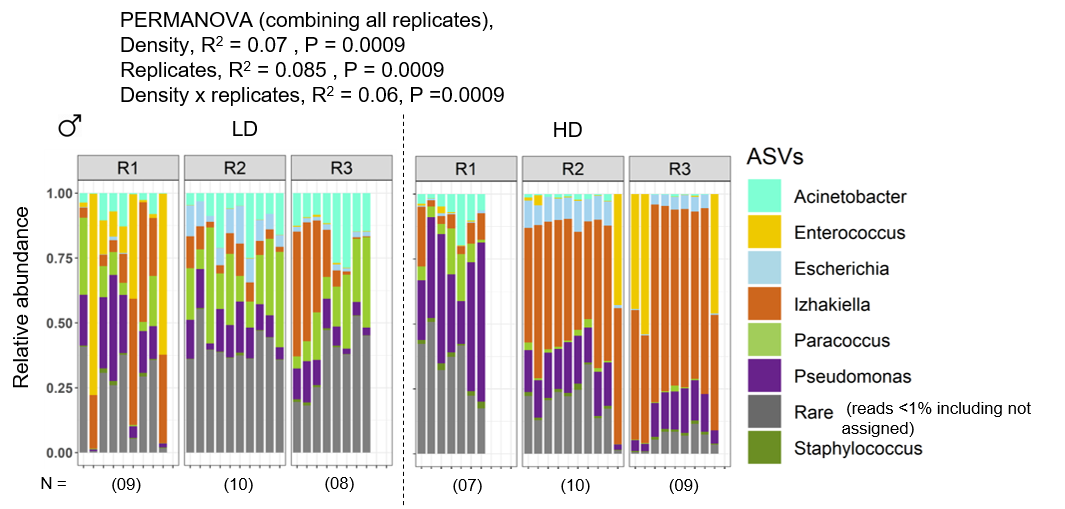

**Fig S15: Larval feces as a key source of beetle microbiome inoculum.** Stacked bar plots show bacterial community composition in the feces of (A) larvae and (B) adults from Experiment 6, as they develop and age. (C) Bacterial community composition in female beetles from different flour treatments (L=larvae, A=adult, indicates source of feces added to flour), from Experiment 7. (D) Flour bacterial community composition on adding different amount of larval feces (% feces by weight indicated on top of each panel), in Experiment 8. (E) Microbiomes of female beetles reared in different flour treatments shown in panel D.

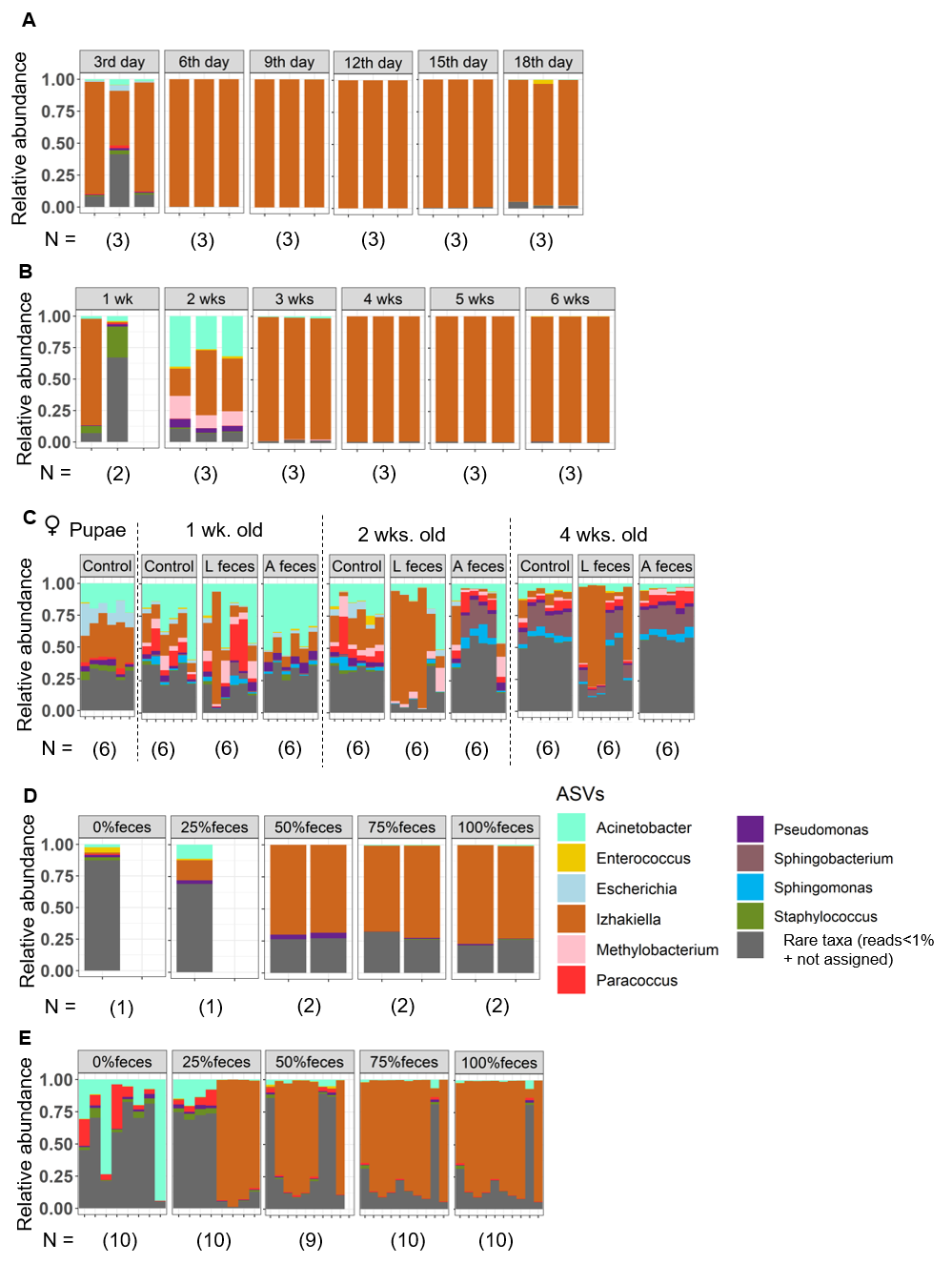

**Fig S16: Impact of quinones on microbiome.** (A) Results of Experiment 10. Flour bacterial communities with different bacterial inoculum (lyophilized bacteria or feces), either with or without added quinones (T0 = initial time point; T4 or T4Q = 16^th^ day time point without or with quinones). (B) Mean (±se) normalized total bacterial load in flour and in beetles reared in the same flour, when relevant (Experiment 7). For flour samples, n = 2 replicates/treatment/time; and for beetle samples, n = 3 females/treatment/time. (C) Microbial community composition of female beetles with or without stink glands, generated using RNAi (Experiment 11). Sample sizes for microbiome analysis are given in parentheses under bars.

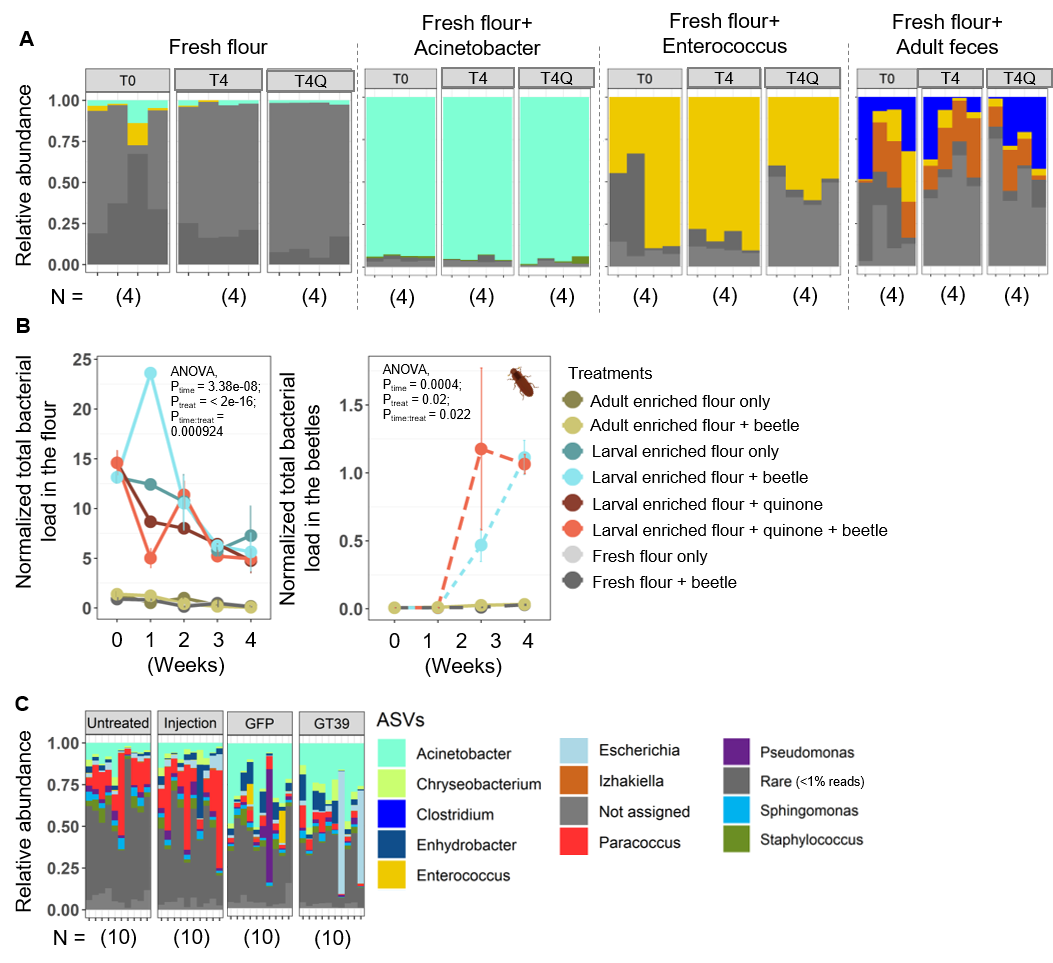

**Fig S17: Impact of flour change on the beetle microbiome.** (A–D) Results of Experiment 12. Stacked bar plots show bacterial community composition in (A) flour, and (B) beetles sampled from a discrete-generation stock population. (C) Flour sampled from a continuous-generation stock population. ‘Cycle’ indicates generations; flour was sampled every 5 days from the time of population initiation; beetles were sampled at different life stages as indicated by schematics above each panel. (D) The relationship between mean total bacterial load and relative abundance of *Izhakiella* in flour samples for samples shown in panels A and C (ANOVA, load ~ relative abundance of *Izhakiella* (rel) x population (pop); P_rel_ <0.0001, P_pop_ = 0.7, P_rel x pop_ = 0.9).

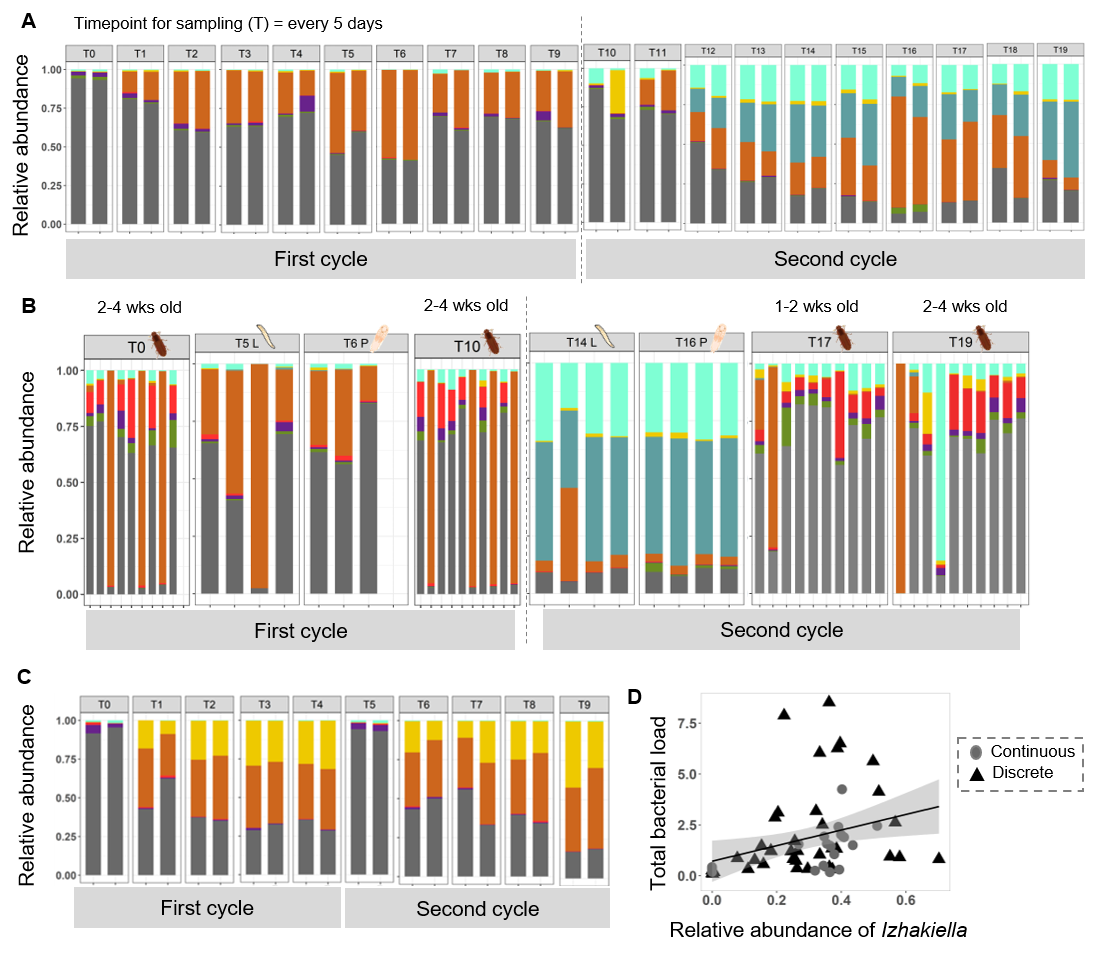
